# Optical pooled screening of a bacterial transposon mutagenesis library

**DOI:** 10.64898/2026.07.07.736948

**Authors:** Oscar Broström, Praneeth Karempudi, Elias Amselem, Maria Tenje, Johan Elf

## Abstract

Transposon mutagenesis is a powerful method to create deep libraries of genetically diverse cells. However, it has previously not been possible to analyze transposon libraries with respect to complex phenotypes as characterized by intracellular spatial dynamics. Here, we use optical pooled screening to characterize a transposon mutagenesis library via live cell single-particle imaging. The library is analyzed in real time, which allows us to use an optical tweezer to isolate cells with interesting phenotypes. We used the method to identify mutants with perturbations in replication initiation control in *Escherichia coli*, but it can in principle be used to identify genetic elements associated with any type of complex or dynamic single-cell phenotype.

## Introduction

Transposon-mediated mutagenesis has become a standard method to probe genotype-phenotype relationships^1^. By inserting themselves into the genome, they disrupt the local genomic region, potentially leading to an altered phenotype. Since the insertions are random, the position of each transposon in the genome must be determined to establish links between genotype and phenotype. With the advent of next-generation sequencing technologies, it became possible to perform large-scale genetic screens of transposon mutagenesis libraries^2–5^. Collectively, these methods are referred to as transposon insertion sequencing (TIS). Originally, TIS was used to find essential genes by utilizing that insertions in essential genes decrease or abolish growth, meaning that few reads in a gene hint at its essentiality. Since then, TIS has been combined with other phenotypic measurements, such as total cell fluorescence, thereby allowing fluorescence-activated cell sorting (FACS) of individual mutants based on intensities^6^. TIS has also been combined with droplet microfluidics to sort out individual mutants for further characterization^7^. Although the extensions of TIS are powerful, they are limited in the complexity of phenotypes that can be studied and for how long.

Microscopy-based techniques lend themselves well to studying more diverse phenotypes, including molecular localization and intracellular dynamics over extended periods of time. The mother machine microfluidic device^8^ has enabled such measurements in bacteria, leading to discoveries in cell cycle control^9–13^, starvation dynamics^14^, antibiotic resistance^15,16^, and bacterial stress response^17^, to name a few. These studies, however, only used one or a few strains per experiment. To increase throughput, we previously developed optical pooled screening^18–21^ (OPS) to be able to screen live-cell strain libraries. OPS starts with a phenotyping stage, followed by an *in situ* genotyping stage where strains are identified using a strain-specific barcode. To read out the barcode, cells have to be fixed, meaning that it is not possible to further characterize strains of interest. Luro et al.^22^ developed the SIFT (single-cell isolation following time-lapse imaging) method to screen non-barcoded strain libraries for dynamic phenotypes. SIFT uses an optical tweezer to isolate cells displaying an outlier phenotype from a strain library grown in a custom-designed microfluidic device. Their device consists of two polydimethylsiloxane (PDMS) layers, which enables compartmentalization, with one clean collection side and one side containing the pooled library where the compartments are separated by pneumatically controlled valves^23^. An elaborate cleaning protocol is required to maintain sterility. For phenotyping, they measure the total fluorescence signal from freely diffusing fluorescent molecules. In this work, we have further developed SIFT by changing the design of the microfluidic device so that multilayer fabrication, which is prone to misalignment and faulty bonding between the layers^24^, and extensive cleaning are not needed; adding single-particle detection sensitivity; and introducing real-time image analysis (segmentation and intracellular foci localization) to increase analysis turnover. We combined this with transposon insertion sequencing, a combination that we will refer to as SIFT-TIS.

We applied SIFT-TIS to search for regulatory elements involved in replication initiation control in *Escherichia coli*. Such regulatory elements are predicted to be involved in cycling the initiator protein DnaA between its active ATP-bound and inactive ADP-bound forms^13^. Known regulatory elements are either expressed gene products or loci in intergenic regions (for details, see Broström^25^ and references therein). Presumably, additional elements fit into these categories as well. Transposon mutagenesis allows for disruptions across the genome, thereby covering both categories. However, most insertions should not be of interest as they would not affect replication. SIFT-TIS therefore lends itself well to study this biological process, as it allows the isolation of transposon mutants displaying rare phenotypes detected at the level of single particles in living cells.

## Results

### Method overview

The basis of our optical pooled screen is a transposon mutagenesis library. The library was created using a conjugative suicide vector that delivers the *magellan6* transposon^2^ to a recipient strain, referred to as the reference strain. The reference strain lacks the flagellar transcription factor *fliA* to facilitate single-cell isolation, and has the fluorescent protein YPet-DnaN fused to the β-clamp of the *Escherichia coli* replisome (YPet-DnaN), which produces foci during ongoing replication (Fig. 1a and Extended Data Fig. 1). The library contains mutants with 31,239 unique transposon insertions distributed in 3,691 genes and 2,098 intergenic regions (Supplementary Tables 1 and 2). To monitor phenotypes, the library is loaded into a mother machine-like microfluidic device^8,26^ designed to allow us to isolate single cells. The device consists of 14 individual segments, each containing 840 trapping channels, referred to as traps (totaling 11,760 traps, but we use two segments at most; see Supplementary Note 1 for details) (Fig. 1b). Upon loading, each trap may contain several different mutants, but after a few hours, each trap will only contain a single mutant. This happens because cells can only enter and exit in one end of the trap, leaving only descendants of the cell that first entered. Phenotypes of individual mutants can therefore be probed using time-lapse phase-contrast and fluorescence microscopy. To process both types of microscopy images, we developed an image analysis pipeline that performs cell segmentation and foci detection in real time (Fig. 1c). We are interested in mutants that deviate from wild-type in terms of replication initiation (from now on initiation) phenotype.

**Fig. 1.**
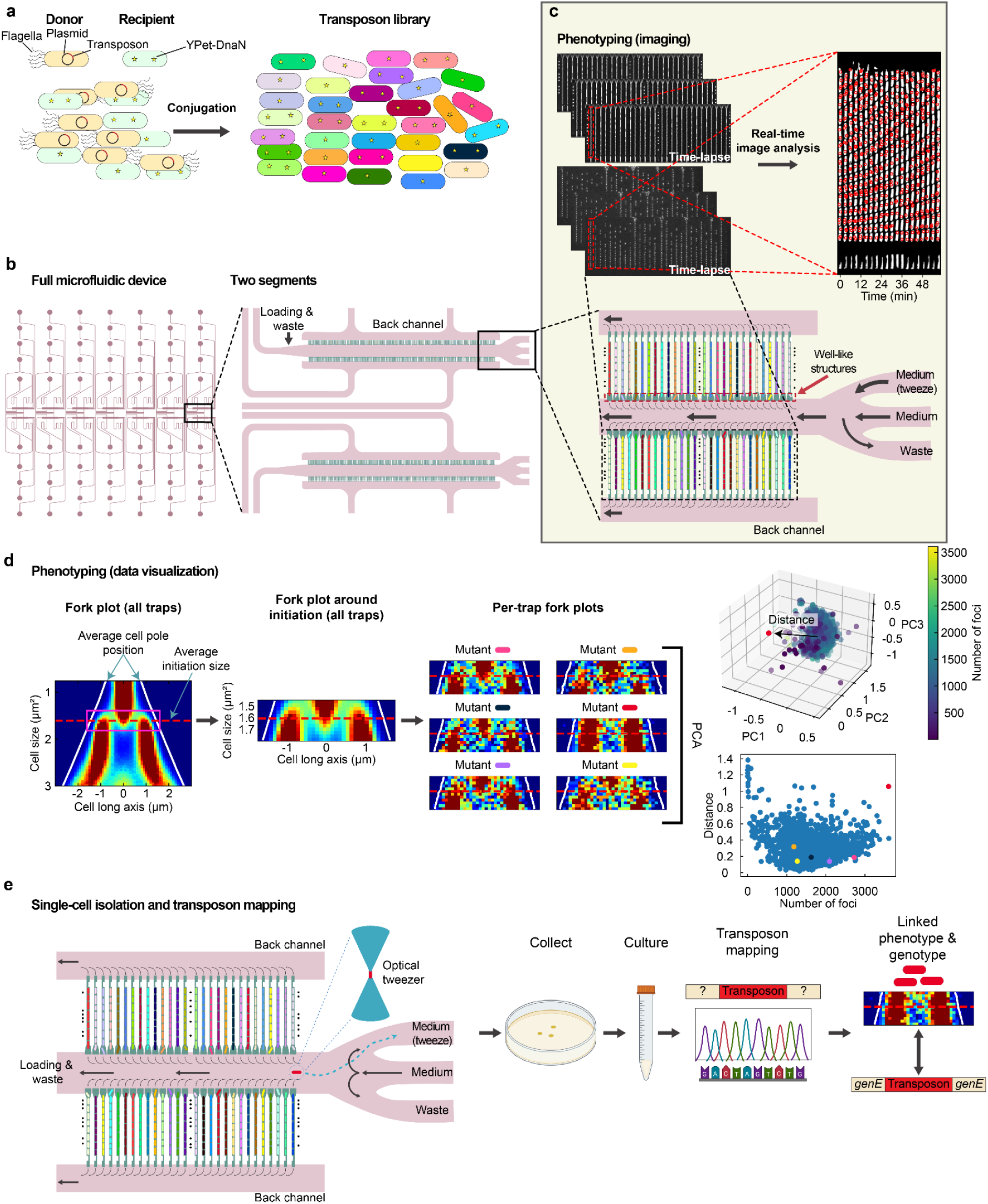
Experimental setup and workflow. **a**, Transposon library construction. A donor strain carrying a conjugative suicide vector is used to deliver the *magellan6* transposon to a recipient strain. The recipient strain has the replisome fluorescently labeled (YPet-DnaN) and the flagellar transcription factor *fliA* knocked out. **b,** Design of the microfluidic device. Left: Full design consisting of 14 independent segments. Right: Zoom-in on the trapping region of two segments. **c,** Phenotyping through microscopy. Bottom: Zoom-in of the trapping region close to the branching channels in one of the segments. Black arrows indicate flow directions and their thickness indicates flow rates (not to scale). Different library mutants are indicated in different colors. Top-left: Time-lapse phase-contrast and fluorescence images. Top-right: Kymograph of segmentation masks and YPet-DnaN foci locations (red circles) of a single trap in increments of 3 min. **d,** Finding deviating initiation phenotypes. Phenotypic data is visualized in a fork plot, which shows the distribution of YPet-DnaN foci along the cell long axis at different cell sizes (µm^2^). The red dashed line indicates the average initiation size and white solid lines indicate the average cell pole positions. Left: Fork plot of all traps in a screen. The box in magenta is a zoomed-in region around the average initiation size for all cells (see Methods for details). The region is used to create initiation fork plots for each trap (mutant). Principal component analysis (PCA) is performed on the fork plots of each trap (see Methods for details), where the first three principal components are visualized (the red point indicates the principal components of an example trap). Deviating phenotypes are classified based on their Euclidean distance from the average point of all traps. The number of detected foci within the initiation region is used to disregard traps with little data for single-cell isolation. **e,** Single-cell isolation and genotype-phenotype linking. Cells are moved to the Medium (tweezer) flow line using an optical tweezer and flushed out of the device. Cells from different traps are collected on separate agar plates. Each colony is cultured and stored so that the transposon location can be mapped. The panel was created using assets from BioRender.

These deviations include changes or increased variability in initiation size, a parameter that normally has a narrow distribution^27^. To estimate these properties, we visualize distributions of the replisome locations on the cell long axis as a function of cell size, called fork plots (Fig 1d). Each branch in a fork plot corresponds to one round of replication and the start of new branches corresponds to initiation. To detect deviations from the wild-type initiation phenotype, we assume that most transposon insertions do not affect initiation control and therefore have the same phenotype as the reference strain, which is similar to wild-type (Extended Data Fig. 1), and focus on a region in the fork plot around ± 11% of the initiation size of all cells in an experiment. We refer to such a fork plot as an initiation fork plot (see Methods for definition). To classify phenotypes in individual traps, the distribution in the initiation region is subjected to principal component analysis (PCA) (see Methods section for details). When observing the first three principal components of each trap, there is a main cluster with a few outliers. The main cluster corresponds to mutants where the transposon insertion does not affect initiation, while the outliers have perturbed initiation. We determine outliers based on the Euclidean distance from the cluster center. If the outliers’ initiation fork plots contain more than 300 foci, we isolate cells from them using an optical tweezer (Fig. 1e and Supplementary Video 1). In Supplementary Fig. 1 we show that we can isolate cells from the trap we want. Collected cells are stored to have the locations of their transposon mapped to link genotype and phenotype, and for further characterization.

### Detection of deviating phenotypes

We first validated whether PCA based on data from individual traps could differentiate strains known to exhibit deviating initiation phenotypes compared to the reference strain. We used strains with knockouts in systems that regulate the nucleotide-bound state of the initiator protein DnaA^28^. The knockouts result in a shift in initiation size; single knockouts of the two DnaA-reactivating sequences (DARS) result in larger initiation sizes, while a knockout of the locus *datA* results in a smaller initiation size, as *datA* is part of the *datA*-dependent DnaA hydrolysis process that deactivates DnaA^13^. We subsequently loaded the knockout and reference strains in mother machine-like microfluidic devices^26^, keeping them separate from one another (this setup will from now on be called arrayed screen). Individually, the strains showed similar phenotypes as we had previously observed^13^ (Extended Data Fig. 2). To see how the knockout strains deviated from the reference, we analyzed replisome foci in a size range around the reference’s average initiation size (see Methods for details) (Fig. 2a). To visualize the initiation fork plot of a typical trap of each knockout, we downsampled the full distributions to match with the number of foci from the median trap (excluding traps with no detected foci) (Fig. 2b). Subsequent PCA of data from individual traps for each knockout resulted in four separate clusters with barely any overlap. Only one trap out of 319 ΔDARS2 traps fell within a convex hull placed around the reference cluster (Fig. 2c). In this screen, the reference strain represents the cluster that would be seen for the transposon mutant library (Fig. 1d), while the DnaA-ATP/ADP regulatory mutants should be regarded as outliers. To test whether this was the case, we calculated the distance of the data point from each trap in the screen from the center of the reference cluster (Fig. 2d). The result reflects the degree of phenotypic deviation from the reference strain, as ΔDARS1 has the most severe phenotype and is farthest away, followed by Δ*datA* and then ΔDARS2. In a replicate experiment (Supplementary Fig. 2) there were more traps from the knockdowns that fell within the convex hull, but there was still separation between the knockouts (see Supplementary Note 2 for details). We conclude that PCA can be used to detect deviating phenotypes similar to those resulting from ΔDARS or Δ*datA*.

**Fig. 2.**
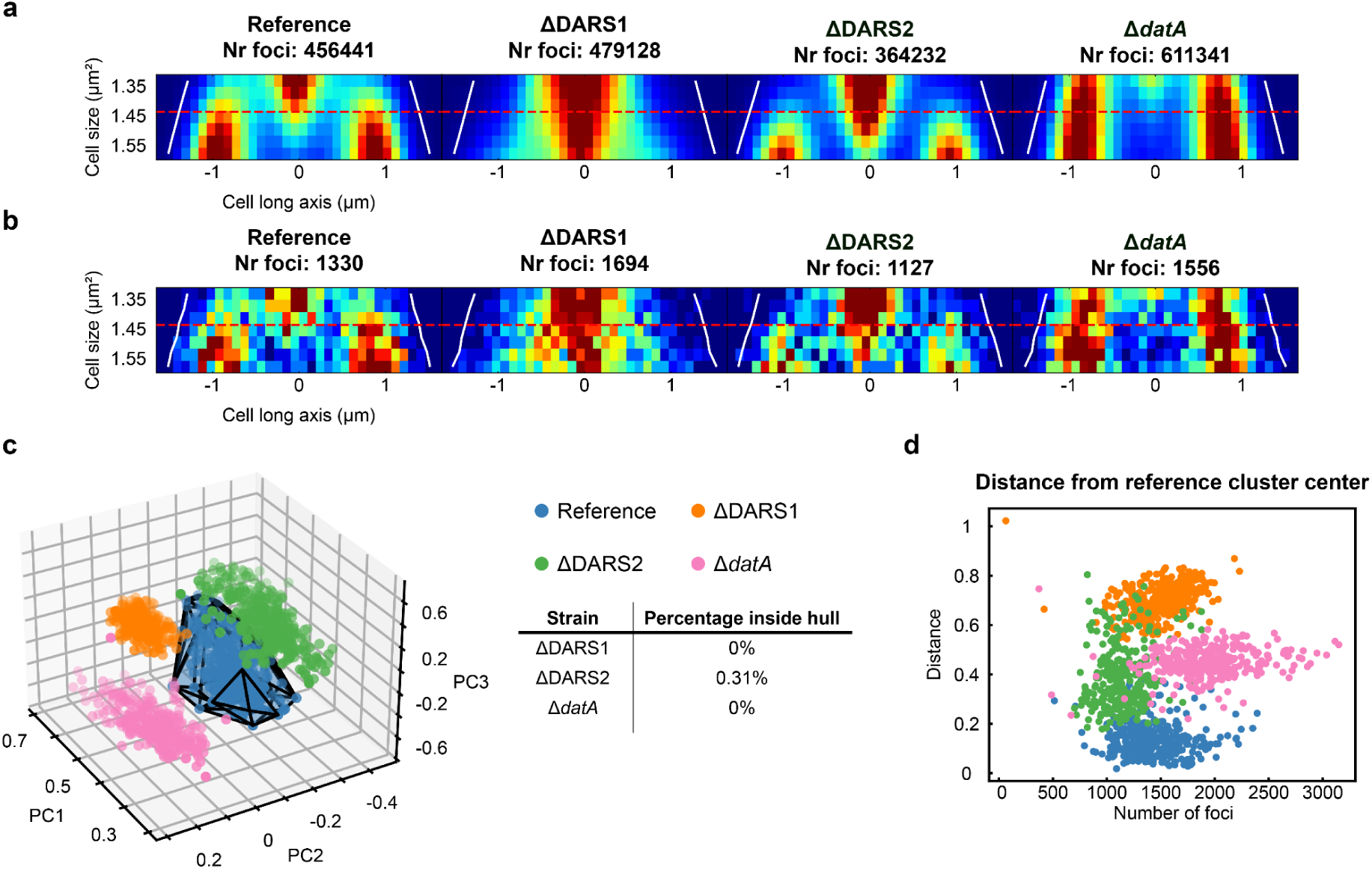
Evaluation of detecting deviating phenotypes. **a**, Initiation fork plots for all traps of DnaA-ATP/ADP regulatory knockouts and the reference strain grown separately from each other. The size range is set by the average initiation size of the reference strain (see Methods for details). The red dashed line in all plots is the average initiation size for the reference strain. **b,** Downsampled fork plots using the same initiation size range as in **(a)**. **c,** PCA of the initiation fork plots for each trap. A convex hull envelopes the reference strain traps (blue). The table indicates the percentage of traps of the DnaA-ATP/ADP knockouts (ΔDARS1: orange; ΔDARS2: green; Δ*datA*: pink) that fall within the convex hull. **d,** Euclidean distance for each trap from the center of the principal components of all reference traps.

### Isolation of mutants with deviating phenotypes from small-scale libraries

To test if we could identify and isolate relevant mutants in a pooled library, we mixed the strains from Fig. 2 so that the reference is overrepresented, leading to the fork plot for the whole pool being similar to the reference (Fig. 3a and Extended Data Fig. 3a). This should mimic what is expected for the transposon mutant library. Subsequent per-trap distance analysis revealed 30 outlier traps from which we chose six traps to isolate cells from (Fig. 3b). These traps were chosen based on their visual similarity (Fig. 3c and Extended Data Fig. 3b) to Fig. 2b so that all three knockouts were isolated. To figure out which trap contained which knockout, we performed colony PCR over the regions of the deletion on each isolated colony grown from single cells (Extended Data Fig. 3c) and used the difference in gel electrophoretic band size to know what deletion was in each trap (Extended Data Fig. 3d–f). The established genotype-phenotype links agreed with our prediction (Fig. 3c). Supplementary Fig. 3 shows a replicate where seven out of eight traps were correctly predicted (see Supplementary Note 3 for details).

**Fig. 3.**
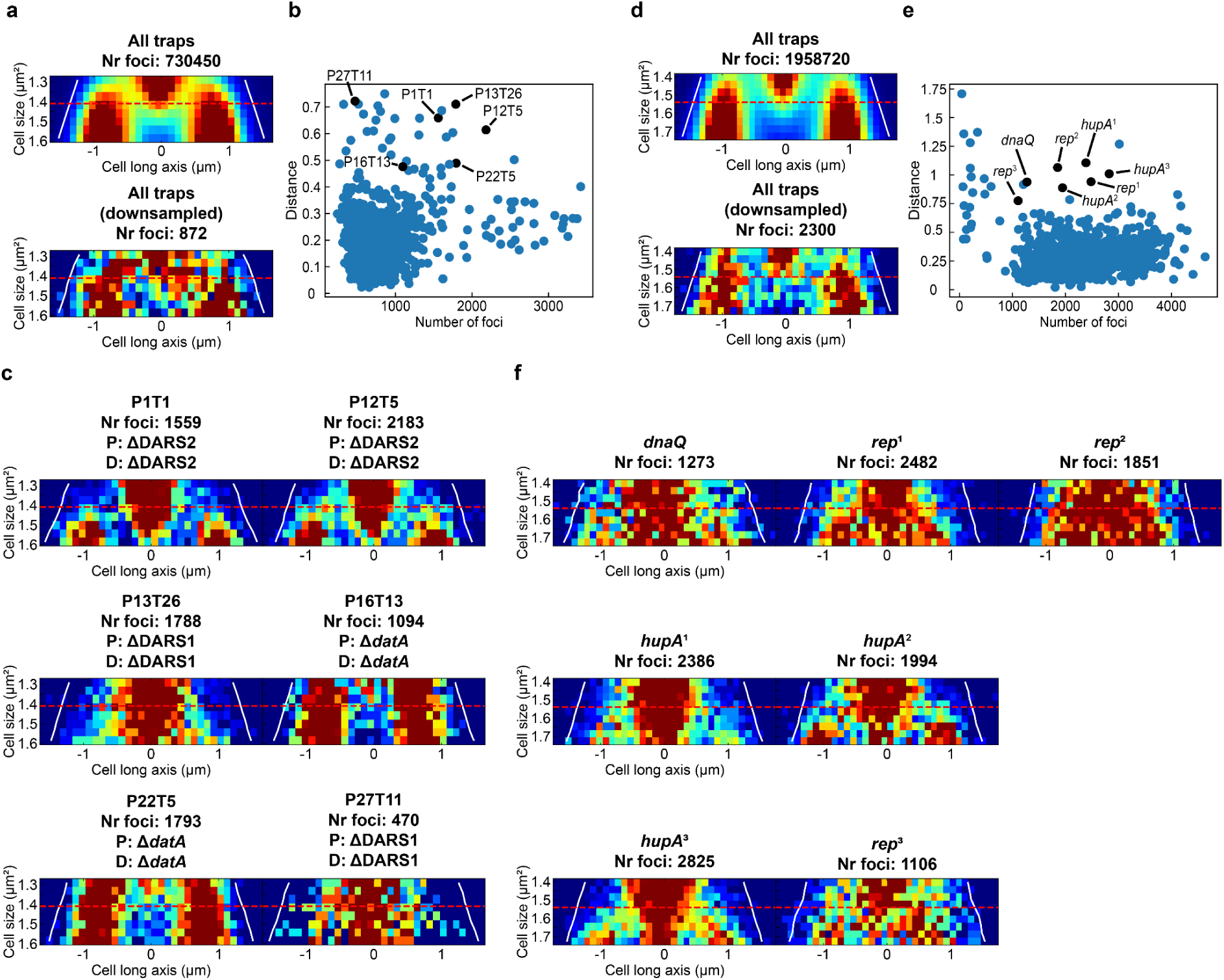
Small-scale single-cell isolation protocol validation. **a–c**, Pooled screen with the reference strain and DnaA-ATP/ADP regulatory knockouts (see Methods for pooling details). **a,** Top: Initiation fork plots for all traps. Bottom: Downsampled initiation fork plot. **b,** Euclidean distance from the PCA cluster center for all traps. Black points indicate the traps from which cells were isolated. **c,** Initiation fork plots for the traps where cells were isolated from. The name of each trap and the number of foci are indicated above each plot. The predicted genotype based on each fork plot’s similarity to Fig. 2b is indicated with P, and the determined genotype by colony PCR is indicated with D. **d–f,** Pooled screen with a CRISPRi library against cell cycle-related genes^20^ (see Methods for pooling details). **d,** Same structure as **(a)**. **e,** Same structure as **(b)**. **f,** Initiation fork plots for traps where cells were isolated from. The number of detected foci and the genes targeted by the respective sgRNAs are indicated above each plot. Lines as in Fig. 2a.

As a second control to confirm that we could find relevant mutants, we used a previously established CRISPRi library targeting genes involved in cell cycle control^20^. A 40-knockdown subpool of the library was mixed (see Methods for details) with the CRISPRi library reference strain so that the reference was overrepresented and subsequently loaded into the microfluidic device for phenotyping (Fig. 3d and Extended Data Fig. 4a). Based on the per-trap distance analysis, eight different knockdowns were isolated (Fig. 3e). These knockdowns varied in their initiation phenotype (Fig. 3f and Extended Data Fig. 4b). The genotype-phenotype link was established by, for each isolated colony (Extended Data Fig. 4c), sequencing the barcode and sgRNA region of the plasmid that each knockdown carried. In this screen (see replicate in Supplementary Fig. 4 and Supplementary Note 3), the isolated knockdowns were all classified as having an effect on initiation control or replication in general (*dnaQ*, *rep*, and *hupA*). *rep* and *hupA* knockdowns were isolated from multiple traps, likely because of their pronounced phenotype, but also that, for this particular screen, more traps happened to be populated by these knockdowns. Overall, we are able to identify and isolate mutants with deviating initiation phenotypes from two small-scale pooled libraries.

### Identification of mutants from the transposon library

To identify initiation control mutants from the transposon library, we have performed five independent screens, with one highlighted in Fig. 4 (the rest are shown in Extended Data Fig. 5). Pooling the data from all traps results in a reference-like fork plot (Fig. 4a). In the screen shown in Fig. 4, we found 14 traps with deviating phenotypes (Fig. 4b). We picked cells from 13 of those traps (Extended Data Fig. 6a) and successfully established a genotype-phenotype link for eight of them. Links were not established either because cell recovery was unsuccessful or because the transposase gene remained in the mutant (see Supplementary Note 4 for details). For traps with successful linking, the phenotypes varied mainly in initiation size (Fig. 4c). We found transposon insertions in seven different genes with various functions (Fig. 4d). Firstly, there was one in *hupB*, which codes for the β-subunit of the nucleoid-associated protein HU^29^. Inhibition of *hupB* expression has previously been shown to result in a larger initiation size^20^. We see this as a positive control that we can identify relevant transposon mutants. Secondly, there were two separate transposon insertions in *dusB*. *dusB* is located in a bicistronic operon upstream of *fis*^30^. Fis is known to affect initiation through DARS2^31^ and through binding to *oriC*^32^. Inhibition of *fis* expression also results in a larger initiation size^20^. This could explain the shift in initiation size observed in our *dusB* mutants. Lastly, the five other genes with transposon insertions (*wzzE*, *hyfJ*, *rbsK*, *rffG*, and *lpoA*) have no known connection to initiation control. It is therefore warranted to further investigate these and other genes found in our other screens (see Supplementary Table 3 for a list of all genes with a genotype-phenotype link).

**Fig. 4.**
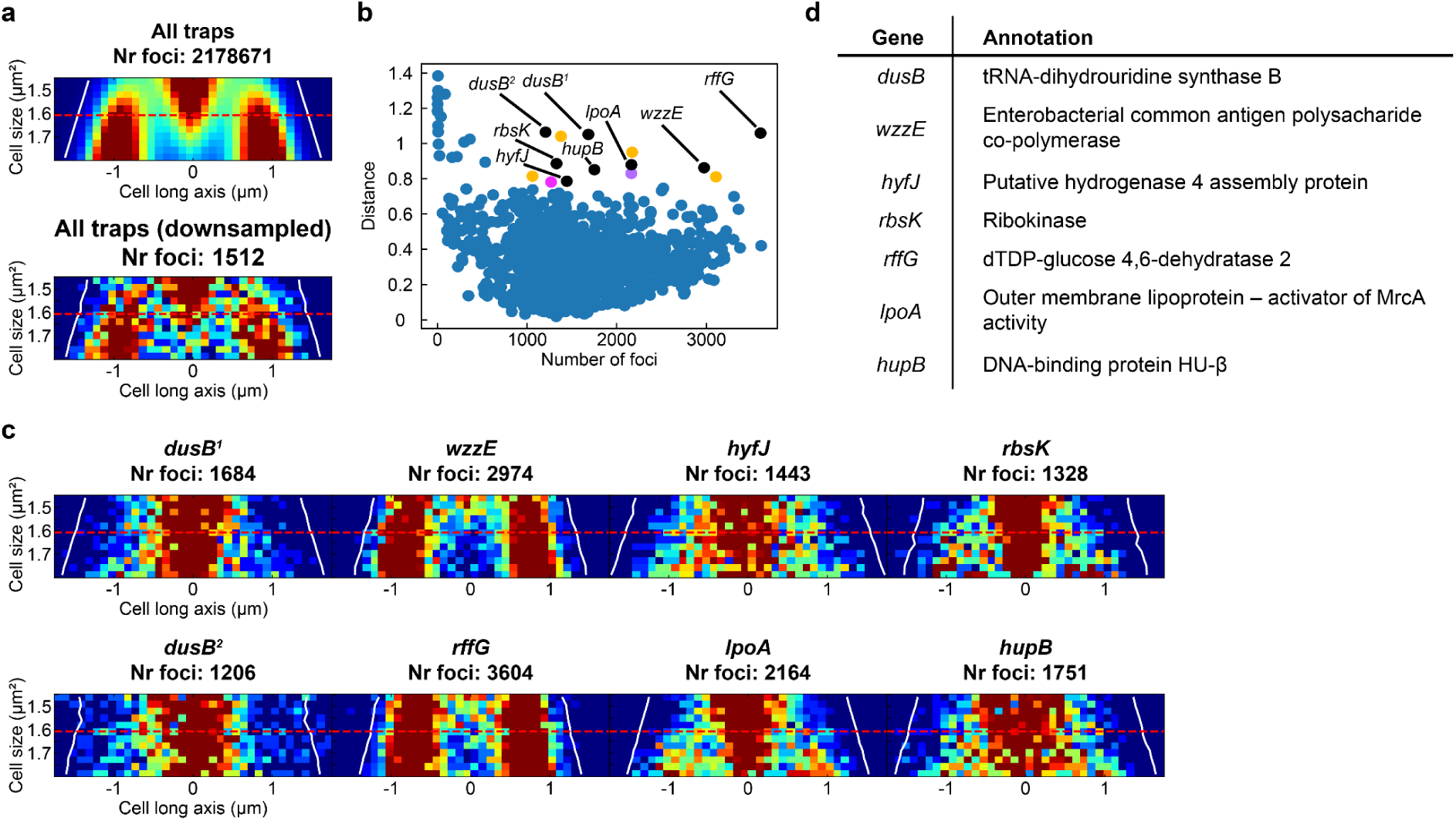
Example pooled transposon library screen. **a**, Top: Initiation fork plot for all traps. Bottom: Downsampled fork plot. **b,** Euclidean distance from the center of all traps. Black: traps with successful genotype-phenotype linking; orange: traps where mutants were not successfully recovered; purple: a trap with successful recovery, but the mutant had the gene for the transposase MarC9; magenta: a trap that was within the threshold used for distance and the number of foci, but no cells were picked from it. **c,** Initiation fork plots for the traps with successful recovery and transposon mapping. The two *dusB* mutants have different transposon insertion sites. **d,** EcoCyc (https://ecocyc.org/) annotations for the genes where transposons were mapped to.

We further investigated 12 out of the total 35 outliers (found in all five pooled replicate transposon screens) with established genotype-phenotype links by running arrayed screens that include over one hundred traps for each of these mutants (Extended Data Fig. 7, Supplementary Fig. 5, and Supplementary Fig. 6). Fig. 5 showcases five of the mutants shown in Fig. 4. Four of these mutants show a similar phenotype (*dusB*, *hupB*, *rffG*, and *wzzE*) when comparing the single trap from the pooled screen (Fig. 5a) and the arrayed screens (Fig. 5b), also at the level of single traps (Supplementary Fig. 6b and 6c). The results for *hupB* and *dusB* are expected based on their direct or indirect role (*dusB* through *fis*) in initiation control. *rffG* and *wzzE*, on the other hand, are new potential genes to investigate further using other methods to probe potential mechanisms. The one gene in Fig. 5 where the results are not consistent between the screens is *hyfJ*. In total there were five mutants with consistent results between the two types of screens and seven mutants with inconsistent results (Extended Data Fig. 7 and Supplementary Fig. 5). On the level of single traps, the *hyfJ* mutant displayed phenotypic heterogeneity (Supplementary Fig. 6b), while the degree of heterogeneity varied for other mutants (Supplementary Figs. 6 and 7; see Supplementary Note 5 for discussion). We reason that the inconsistencies could have arisen from mutations that occurred at some point between single-cell isolation and testing the mutants separately. To check this, we performed whole-genome sequencing. Out of the sequenced mutants, only the *hyfJ* mutant had acquired an additional mutation in the gene *pntA*, but it is unlikely that this mutation is the cause of the discrepancy, as the proteins HyfJ and PntA are involved in different biological processes (see Supplementary Note 6 for details). Another possibility is that a few traps appear deviating in the transposon screen simply because they are undersampled in the single-trap analysis, *i.e.*, that a clear separation from the cluster in the PCA analysis is not a sufficient criterion compared to being different in an arrayed screen with much more data.

**Fig. 5.**
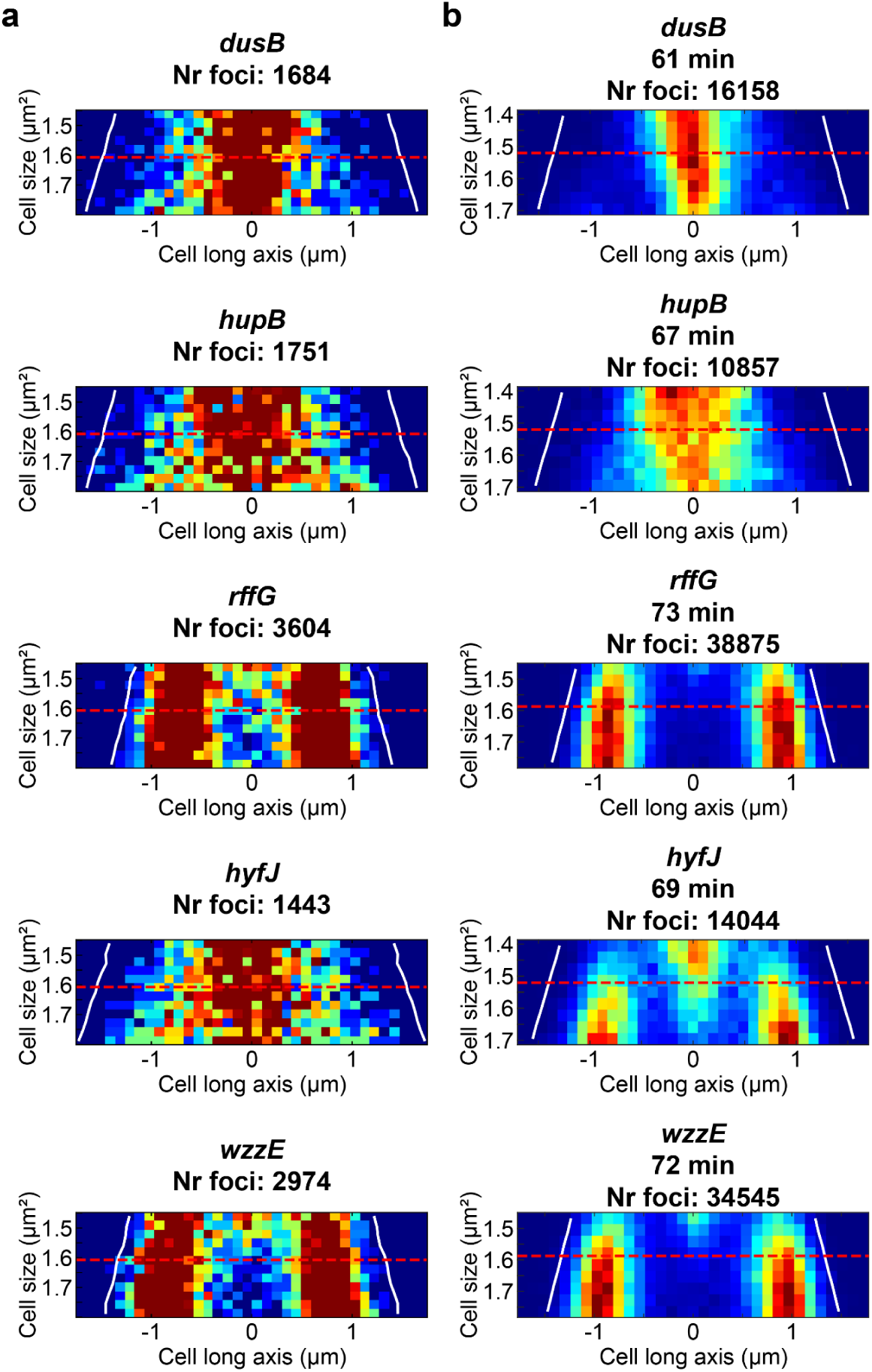
Comparing pooled and arrayed screens of individual transposon mutants. **a**, Initiation fork plots for pooled library screens (same as in Fig. 4c). **b,** Initiation fork plots for multiple traps of single mutants. The red dashed line corresponds to the average initiation size of the reference strain run in the same screen (see Supplementary Fig. 5). The average generation time of each mutant is indicated above each plot.

## Discussion

Transposon insertion sequencing has enabled many discoveries, for example, conditionally essential genes and virulence and host adaptation of pathogens, to name a few^1^. However, transposon mutagenesis has previously not been used to study complex phenotypes related to intracellular dynamics. We have developed a method to screen a transposon mutagenesis library for defects in initiation control, allowing us to identify mutants with deviating phenotypes and trace them back to their underlying genetic causes. Using this approach, we identified 35 such cases involving different genes and intergenic regions (Fig. 4, Extended Data Fig. 5, and Supplementary Table 3). For 12 of these, we performed additional imaging with higher sampling to confirm the phenotypes (Fig. 5, Extended Data Fig. 7, and Supplementary Fig. 5). Five mutants (*dusB*, *hupB*, *rffG*, *wzzE*, and *pabA*) showed consistent behavior across both pooled and arrayed screens. Further discussion on each mutant can be found in the Supplementary Note 6. In particular, the genes *rffG* and *wzzE* should be investigated further in regard to their role in initiation control.

An advantage of mother machine-type phenotyping is that individual strains do not compete with one another, allowing the diversity of the library to be maintained over extended periods despite differences in growth rate. In the present setup, we did not, however, monitor growth rates at the level of individual traps, as the imaging frequency (every 3 minutes) does not allow single-cell tracking in our growth conditions. This limitation is partly due to the current speed of real-time image analysis. In practice, growth defects can confound the analysis, as changes indirectly affect the measured initiation behavior, making it difficult to distinguish true defects from general physiological impairments. Incorporating growth rate measurements alongside initiation would allow us to separate these effects.

At present, we monitor approximately 1,680 traps per screen. This number could be substantially increased by minor changes in fluidic routing (see Supplementary Note 1 for details). However, the number of cells that can be isolated per screen would not scale proportionally. Consequently, selection thresholds for targeting cells would need to become more stringent in order to prioritize the most informative phenotypes if we scale up to monitor more mutants. More generally, the total number of cells analyzed will depend strongly on the specificity and frequency of the phenotype of interest. This could be determined by real-time statistical analysis to figure out how many outliers meet specific criteria.

A general limitation of transposon mutagenesis libraries is that mutants with severe fitness defects will not be in the library. A promising development is inducible transposon mutagenesis^33^, where libraries could be generated in the microfluidic device. This would allow for phenotyping of mutants with lethal transposon insertions. It would, however, require single-cell whole-genome amplification to map the transposon insertion since the mutant will eventually die.

SIFT-TIS allows for the identification of connections between rare genotypes and complex dynamic phenotypes, any genetic perturbation related to a transposon insertion, and any phenotype that can be detected by super-resolved time-lapse microscopy. Only changes to the phenotypic analysis would be necessary to investigate phenotypes of other transposon mutagenesis libraries.

## Methods

### Strains, cloning, and media

Strain construction was done in LB. Supplements were added to maintain plasmids or to keep some strains viable. All strains used in this study are listed in Supplementary Table 4. All oligos can be found in Supplementary Table 5. In the fluidics experiments, cells were grown in M9, 100 μM CaCl_2_, 2 mM MgSO_4_, F-108 Pluronic^®^ (425 µg/ml or 51 µg/ml, see Supplementary Table 6 for details), 0.4% succinate (w/v), 1x RPMI 1640 amino acids. For the CRISPRi library, the medium was supplemented with 0.1 mM uracil and 50 µg/ml kanamycin.

#### Transposon library reference strain & DnaA-ATP/ADP regulatory knockouts

Generalized P1 transduction^34^ was used to transfer genetic constructs. For the transposon library reference strain, the genetic constructs *kan-ypet-dnaN* and Δ*fliA*::*kan* were transferred sequentially. In between transferring each construct, the kanamycin resistance cassette was flipped out using pCP20, leaving *frt* scars. The knockouts of DARS1, DARS2, and *datA* were transferred from previously created constructs^13^.

#### Transposon library construction

To create the transposon mutagenesis library, we used the conjugative suicide vector pMarC9-R6k (Addgene plasmid #89477; http://n2t.net/addgene:89477; RRID:Addgene_89477)^35^, which carries the *magellan6* transposon^2^ and its transposase MarC9. To the plasmid, we introduced a *catsacB* cassette (see Supplementary Note 4 for why this was done). This plasmid, pMarC9-R6k-catsacB, was constructed using In-Fusion cloning (Takara Bio). PCR (always according to the manufacturer’s instructions) was used to linearize pMarC9-R6k and add homology sequences for the plasmid backbone to the *catsacB* cassette. The samples were treated with DpnI, PCR-purified, and then combined together with the In-Fusion master mix. pMarC9-R6k-catsacB was transformed into DH5α carrying the *pir* protein (*pir* is necessary as the plasmid requires it for replication). The plasmid was purified and electroporated into MFD*pir*, a gift from Jean-Marc Ghigo^36^, resulting in our donor strain. These two strains require supplementation of diaminopimelic acid (DAP) to survive and propagate the plasmid.

The transposon library was created by inoculating the reference strain and the donor strain in LB supplemented with 0.3 mM DAP (50 µg/ml kanamycin for the donor strain) and incubating them at 37 °C, 200 rpm, overnight. The cultures were centrifuged, resuspended in LB 0.3 mM DAP, and mixed 1:1 volumetrically. To preheated LA 0.3 mM DAP plates, cellulose nitrate membranes (MilliPore, HAWP02500) were added. Mixed culture was spotted on the membranes, and the plates were incubated for ∼5.5 h at 37 °C. The membranes were transferred to Falcon tubes containing LB. The tubes were vortexed to resuspend the cells in the medium. The cultures were diluted 1:5 and plated on LA 50 µg/ml kanamycin plates and incubated overnight at 37 °C. Plates were harvested (approximately 77,000 colonies) by adding LB, resuspending them with a bacterial spreader, and collecting the resuspended bacteria. The resulting culture was used to make glycerol stocks (six subpools forming one library named Sucrose-), and diluted to be plated onto salt-free LA 5% sucrose plates. The plates were incubated overnight at 37 °C and subsequently harvested the same way as above. This library was stored as glycerol stocks in two subpools, called Sucrose+. Sucrose+ was used for all transposon library screens.

### Microfluidics

#### Microfluidic device design and wafer fabrication

The microfluidic device contains flow channels fabricated at two different heights. The traps where the cells grow are 1.125 µm in width and height and 54 μm in length. The trap design is modified from the open-ended mother machine design^26^ to include a well-like structure at the opening of the trap (Fig. 1C), which helps in accumulating 4–5 cells of the same genotype and makes it easier to isolate cells with the optical tweezer. The rest of the flow channels are 10 μm in height.

The wafer was produced by ConScience AB according to our specifications. The structures are fabricated on a standard 6-inch Si substrate of 525 ± 20 μm thickness. A SiO_2_ layer of thickness 1.125 µm is grown on this substrate using thermal oxidation. The trap design is etched in the SiO_2_ layer using dry reactive ion etching after the trap structures have been defined using e-beam lithography. The larger flow channel design of height 10 μm is fabricated on top of the traps in AZ^®^ 10XT 520cP resist using UV lithography. The larger channels are also curved using resist reflow to allow complete valving of these channels, if needed.

#### Microfluidic device molding

All microfluidic devices were fabricated in polydimethylsiloxane (PDMS, Sylgard 184) bonded to a glass coverslip using air plasma. For the single-cell isolation device, the wafer mold was mounted in a custom aluminum manifold designed to cast devices inside the cavity between its two pieces. Approximately 90 g of PDMS (1:10 base and curing agent) was injected into the cavity using a syringe until the air in the cavity was removed. The aluminum manifold was then placed in an 80 °C oven for 6+ h to cure. For experiments using the same design as Baltekin et al.^26^ (see Supplementary Table 6 for details), the manifold was not used; instead, 45 g of PDMS (1:10 base and curing agent) was poured on the wafer mold. The PDMS was degassed and cured as before.

For all device designs, individual devices were cut out from the PDMS slabs, and inlets and outlets were punched using a 0.5 mm biopsy puncher before the PDMS was bonded to a glass coverslip after plasma treatment using a HPT-200 Benchtop Plasma system (Henniker Plasma) at 50% power for 30 s. Tygon tubing (i.d. = 510 μm, o.d. = 1,524 μm; Saint-Gobain) and custom-made metal pins (23-gauge, 14-mm-long bent in the middle at 90°; New England Small Tubing) were used to interface the bonded chip with a pressure-driven flow controller.

#### Cell loading

From frozen stock, cells were inoculated in the growth medium of the corresponding experiment (see the *Strains, cloning, and media* section) and incubated overnight at 30 °C, 200 rpm. To preheated medium, cells were diluted 1:100, 1:250, 1:500, or 1:1,000 and incubated until the device was ready to be loaded (on the scale of hours).

To maintain flow and for wetting devices, a flow controller was used (OB1 MK3+, Elveflow) for all experiments except the flagella control experiment, where a custom-built pneumatic system was used. The wetting procedure when using devices of the same design as in^26^ was the following (nomenclature as in Figure S1 of Baltekin et al.^26^): back channel (5.1 and 5.2 simultaneously), back waste (6.0), medium in (2.0), and loading/cell waste (2.1 and 2.2 simultaneously). In the arrayed screens two devices were side-by-side, with one device wet at a time. Loading on these devices was done by keeping some pressure on the medium in port while mostly having pressure on the loading ports. Once loaded, the pressure on the medium in port was set to 200 mbar, and the back channel tube was placed 40–60 cm below the other tubes. The temperature of the device was kept at 30 °C by an incubator (Okolab).

On the single-cell isolation device, the order of wetting and pressures was the following: medium (tweezer) port, 700 mbar; medium port, 300 mbar; front waste port, 200 mbar; back channel port, 200 mbar; and loading/cell waste port, 200 mbar. When using two segments, each segment was wet at a time. When loading, the pressure was 80 mbar on the medium (tweezer) port, 70 mbar on the medium port, 0 mbar on the front waste and back channel ports, while the pressure on the loading port was initially set to 50 mbar and subsequently adjusted to not have too many cells flowing past the last traps. With this setup, cells that flow by the last traps flow toward the front waste port, keeping the two medium sources sterile. Once done, the pressures were set to 200 mbar on the medium (tweezer) port and 100 mbar on the medium port, and the back channel tube was placed 40–60 cm below the other tubes.

When isolating fluorescent cells from a pool with non-fluorescent cells (Supplementary Fig. 1), the two strains were mixed 1:10 volumetrically prior to loading. The small-scale DnaA-ATP/ADP regulator library was created by volumetrically mixing the knockouts 1:1:1:40 together with the reference strain. For the CRISPRi library, the chip was wet with medium supplemented with 1 ng/ml anhydrotetracycline (aTc) to induce sgRNA expression. Before loading, the CRISPRi subpool was volumetrically mixed 1:10 with the CRISPRi library reference strain.

One segment of the single-cell isolation device was used for the small-scale library screens. Two segments were used for the transposon library screens.

#### Single-cell isolation

Before cell isolation was started, the pressure on the medium (tweezer) port was turned off. The tubing connecting the microfluidic device and the Falcon tube with medium was then cut with scissors, leaving a 5–10 cm long tube still connected to the device. This tubing was placed onto an LA plate (supplemented with 50 µg/ml kanamycin for the fluorescent cells, and CRISPRi and transposon libraries) to collect liquid.

Cell isolation was done similarly to Karempudi et al.^24^. Briefly, the SLM was configured to shape and focus the IR laser beam to the axial plane where cells are located. A cell is captured and moved by this focal spot. The location of the focal spot was marked with a circle in MicroManager 2.0^37^. A 750 nm shortpass filter (Thorlabs FESH0750) was pushed into the camera path, making the beam invisible on the camera. The trapped cell is moved by moving the stage. A shutter is used to block the laser beam to release the cell. The laser power was approximately 170 mW at the back aperture of the objective.

When isolating cells, the pressure was lowered to approximately 25 mbar on the medium port. Once a cell was trapped, it was dragged towards the tweezing port. When the cell reached the flow line of the channel leading to the medium (tweezer) port, the laser was cut off (Fig. 1e). In general, multiple cells were isolated from each trap to avoid potential issues with cell death from the laser or cells getting stuck. Subsequent rounds of cell isolation from one trap were done as soon as the next cell was at the well-like structure. The time for this to occur varied, as cells had to be in the well-like structure for efficient trapping. While waiting, the pressure on the medium port was changed to 200 mbar. Once a sufficient number of cells were isolated from one trap (typically three), the pressure on the medium port was increased to 500 mbar for 15 min. Subsequently, the tubing at the medium (tweezer) port was disconnected, and the liquid in the tubing was ejected onto the plate using an air-filled 1 ml syringe. To reset the system, a new 5–10 cm tube was wet with medium using a syringe. The tube was connected to the device at the medium (tweezer) port and placed on a fresh LA plate of the same type as before. Once cells had been isolated from all of the desired traps, liquid from a fresh tube was collected on a fresh LA plate (same type as before) to ensure that there was no contamination. The plates were incubated at 37 °C as soon as medium had been ejected onto them. Incubation lasted overnight. For the pooled transposon screens, colonies were inoculated in LB and grown at 37 °C, 200 rpm, for 8 h or overnight and subsequently stored as 20% glycerol stocks at -80 °C.

### Microscopy

#### Optical setup

We used two different optical setups. See Supplementary Table 6 for which experiment was done on which setup.

Optical setup 1: An inverted TI2-E (Nikon) microscope with a CFI Plan Apo lambda 100x, 1.45 NA oil DM objective (Nikon) was used. Images were acquired on a Kinetix (Teledyne Photometrics) camera. Illumination for phase-contrast was through an LED (Nikon), resulting in diascopic illumination. Fluorescence illumination was through a Spectra Gen III (Lumencor) with the TEAL setting set to 6%. For fluorescence illumination, the light was filtered through a filter cube with a FF01-514/3 (Semrock) excitation filter, an ET550/50M (Chroma) emission filter, and a ZT514rdc-UF2 (Chroma) dichroic mirror. The microscope was controlled using MicroManager 1.4.23^37^ with in-house plugins.

Optical setup 2: An inverted TI2-E (Nikon) microscope with a CFI Plan Apo lambda 100x, 1.45 NA oil DM objective (Nikon) was used. Images were acquired on an ORCA-Quest qCMOS (Hamamatsu) camera. Illumination for phase-contrast was through an LED (Nikon), resulting in diascopic illumination. Fluorescence illumination was through a Spectra Gen III (Lumencor) with the TEAL setting set to 6% or 36% (see Supplementary Table 6 for details). For fluorescence illumination, the light was filtered through a filter with a FF01-514/3 (Semrock) excitation filter, an ET550/50M (Chroma) emission filter, and a ZT514rdc-UF2 (Chroma) dichroic mirror. The microscope was controlled through MicroManager 2.0^37^ with appropriate device adapters for the light sources, stage, and camera. When running real-time analysis, the image acquisition was controlled through Pycro-Manager^38^. The optical tweezer setup was custom-built, based on a 1,030 nm 1 W fiber laser (MPBC 2RU-YFL-P-1-1030) and a spatial light modulator (SLM, Hamamatsu X15213-03) for beam focusing and steering. The first-order refraction is isolated and directed to the back focal plane of the objective using a dichroic mirror (Thorlabs DMSP900R). To avoid damaging the camera, a shortpass filter (Thorlabs FESH0750) is inserted into the camera path. Custom-built SLM software was used to change the tweezing plane.

#### Imaging

For the arrayed DnaA-ATP/ADP regulatory knockouts and all pooled screens, images were acquired every 3 min with 30 and 50 ms exposure times for phase-contrast and fluorescence, respectively. Imaging was done for 20 h.

For the Δ*fliA* experiment, phase-contrast images were acquired every 30 s with 50 ms exposure. Fluorescence images were acquired every 2 min with a 300 ms exposure. Imaging was done for 3 h.

When imaging the isolated transposon mutants, phase-contrast images were acquired every minute with a 30 ms exposure. Fluorescence images were acquired every 2 min with a 50 or 300 ms exposure. Imaging was done for 4–7 h (see Supplementary Table 6 for details).

### Image analysis

The amount of data for each experiment in the different figures can be found in Supplementary Table 6.

#### Phenotyping

For the Δ*fliA* experiment and the arrayed transposon mutant screens, images were analyzed using a fully automated image analysis pipeline primarily written in MATLAB (Mathworks, R2024a)^13^. Cell segmentation was done using U-Net-based segmentation^39^. Cell tracking was performed (only done in Fig. 5, Extended Data Figs. 7b and 7d, and Supplementary Fig. 1) using the Baxter algorithm^40^. Fluorescent foci were detected using the wavelet algorithm^41^.

After running the image analysis pipeline, the output was post-processed using custom-written MATLAB scripts and functions^13^. The initiation sizes shown in Fig. 5b, Extended Data Figs. 7b and 7d, and Supplementary Fig. 5 were determined as in Knöppel et al.^13^.

#### Real-time analysis

Real-time image analysis was done using two Python packages. One package (rtseg) did cell and trap segmentation, device barcode detection, fluorescence foci detection, fork plot generation, and trap-wise distance measurements. The second package (rtclient) was used for image acquisition and data visualization.

Cell and trap segmentation was done using a U-Net similar to Kandavali et al.^42^. The U-Net architecture was optimized to segment one image in ∼300 ms. A YOLOv3 network^43^ was used to identify barcodes placed around every 14 traps to aid trap identification. The wavelet algorithm^41^ was re-implemented in PyTorch to, in one image, detect foci in ∼50 ms on a GPU. The foci are then placed in an internal coordinate system for each cell, which took 400–500 ms to determine per image.

The computer controlling the microscope was equipped with an Intel i9-10900K CPU 3.70 GHz processor, an NVIDIA RTX 3090 GPU, and 32 GB RAM. While running, experiments use ∼12 GB of GPU and ∼18 GB of CPU memory, respectively.

#### Per-trap analysis

To get initiation fork plots, the average initiation size is first calculated as in Camsund et al.^20^ (in bulk). Based on the average initiation size, size (y-axis) bins ±11% of it are selected, with one larger size bin added. Along the x-axis (cell long axis coordinate), the range is determined by when the cells are the longest within the y-axis range. PCA is done on a per-bin (pixel) basis on the per-trap initiation fork plots, reducing to three dimensions. In the arrayed DnaA-ATP/ADP regulatory mutant pooled screens, a convex hull was placed around the individual reference strain traps with any number of detected foci in the replicate shown in Fig. 2 and above 300 detected foci in the replicate shown in Supplementary Fig. 2 (see Supplementary Note 2 for details). In pooled screens, the average 3D coordinate is calculated and used for Euclidean distance measurements. In the arrayed screens, the average 3D coordinate of the reference strain is used instead. Outliers are selected using thresholds for the number of foci and distances. When pooling the DnaA-ATP/ADP regulatory mutants, further selection by eye was done to ensure that all mutants were isolated. Cells were never tracked in the per-trap analysis.

### Sequencing

#### Transposon sequencing

Transposon sequencing was done similarly to^44^, but with a few differences. Starter cultures were made by inoculating each previously frozen subpool of the Sucrose- and Sucrose+ libraries from frozen stock in LB. The cultures were incubated at 37 °C, 200 rpm, until OD_600_ = 0.3–0.5. The cultures were centrifuged, the supernatant discarded, and the pellets were resuspended in LB 20% glycerol and stored at -80 °C. Genomic DNA from the starter cultures was prepared by salting out the DNA. The starter cultures were thawed on ice, inoculated in LB, and grown until OD_600_ ≈ 0.1 at 37 °C, 200 rpm. Cultures were harvested by centrifugation and resuspended in TE buffer. Cells were lysed by adding lysozyme (10 mg/ml) followed by incubation at 37 °C for 1–2.5 h. Subsequently, 10% SDS and 20 mg/ml Proteinase K were added, followed by incubation at 55 °C for 3 h. Then, 2 ml of 5 M NaCl was added before vigorously shaking the tubes. Tubes were incubated on ice for 5 min, centrifuged at 4 °C, and the supernatants were transferred to new tubes. Room temperature isopropanol was added (0.6x of the sample volume). Mixing was done by inversion. Tubes were then centrifuged for 10 min. Supernatants were discarded, and the pellets were washed with 70% EtOH. After another centrifugation step and with the supernatants being discarded, the pellets were air-dried at 37 °C. UltraPure H_2_O (Thermo Fisher) was added, and each pellet was allowed to dissolve at 37 °C for 1 h before storing at -20 °C.

The genomic DNA from each subpool of the corresponding library was pooled, then digested with MmeI (NEB), dephosphorylated with Quick CIP (NEB), extracted with phenol:chloroform:isoamyl alcohol (25:24:1, v/v%), and precipitated at -80 °C in 100% pre-chilled EtOH. After centrifugation, the pellet was washed with ice-cold 70% EtOH, air-dried at 37 °C, and resuspended in UltraPure H_2_O (Thermo Fisher). Adapters (0.2 mM) were ligated onto the purified DNA using T4 DNA ligase (400 U/µl, NEB). The adapter pair ADBC-F-INDX1:ADBC-R-INDX1a was used for Sucrose- and ADBC-F-INDX1b_catsacB:ADBC-R-INDX1b_catsacB was used for Sucrose+ (Supplementary Table 5 for adapter sequences). The ligation products were PCR amplified, minimizing the number of PCR cycles. The PCR products were loaded on agarose gels, and the bands with the correct size were excised and purified using the Monarch Gel Extraction Kit (NEB). The purified product was PCR amplified to add Illumina adapters (keeping the number of cycles low). The PCR products were purified, pooled, and sequenced on a MiSeq sequencing system using the v3 sequencing chemistry and paired-end 75 bp read length.

Analysis was done by trimming the reads using Cutadapt^45^ and mapping the reads to the reference genome of the library background in CLC Genomics Workbench (QIAGEN).

Custom-written bash scripts were used to extract the number of unique TA insertion sites and which genes and intergenic regions have transposon insertions in them. Note that the quality of the reverse read was low for both libraries, so only forward reads were used.

#### Whole-genome sequencing

Genomic DNA of the wild-type and library background was prepared using the DNeasy® UltraClean® Microbial Kit (QIAGEN) according to the manufacturer’s instructions. Genomic DNA for the transposon library mutants was the same as for the transposon mapping (see *Genotype identification of isolated cells* and *Transposon sequencing*). The DNA was prepared for sequencing using Illumina DNA Prep according to the manufacturer’s instructions. Index primers are listed in Supplementary Table 5. The cleaned-up DNA from each strain was pooled and sequenced on an iSeq100 sequencing system (Illumina).

Analysis was done by a workflow consisting of adapter trimming by TrimGalore^46^, mapping the reads to a reference using BWA-MEM2^47^, and variant-calling using Pilon^48^. The genome of MG1655 was used as an initial reference genome (accession number U00096.3). Results were refined in CLC Genomics Workbench (QIAGEN).

### Genotype identification of isolated mutants

The genotypes of the isolated DnaA-ATP/ADP regulatory mutants were determined by colony PCR using control primers for each deletion. Positive and negative controls were strains with and without the deletions, respectively. All tweezed colonies were screened, but only with the primers corresponding to the presumed phenotype.

The genotypes of the isolated colonies from the CRISPRi library were determined by doing colony PCR and Sanger sequencing (Eurofins Genomics) over the barcode and sgRNA sequence. Results were aligned to the plasmid sequence in SnapGene (GSL Biotech LLC) or Benchling (Benchling, Inc.), and the barcode and sgRNA sequence were compared with Supplementary Table 5 in Camsund et al.^20^.

For the transposon library, transposon mapping was done similarly to the *Transposon sequencing* section. The differences were that genomic DNA was prepared from stationary-phase cultures, samples were not pooled, adapter sequences and PCR primers were different (Supplementary Table 5), in some of these experiments the adapter concentration was 20 µM, the number of PCR cycles was different, and sequencing was done through Sanger sequencing (Eurofins Genomics). Only one colony from each trap was used for mapping. For some isolated mutants, PCR over the transposon insertion sites was done to check that all colonies collected on one plate had the same insertion site.

### Use of large language models

Perplexity AI (https://www.perplexity.ai/) was used to assist in the development of the real-time analysis pipeline, developing the bash scripts used for TIS analysis, and developing the protocol for salting out DNA.

## Acknowledgements

We would like to thank Jimmy Larsson for input on protocol development and microfluidic device design, Johan Öhman for assistance with device design, Dvir Schirman for help with NGS, Beer Chakra Sen for help with NGS and advice on strain construction, David Fange for suggesting the CRISPRi library control and careful proof-reading of the manuscript, Daniel Jones for constructing the Δ*fliA*::*kanR* construct and providing the reference barcode sequence for the CRIPSRi library, Anna Knöppel for strain construction and transposon library construction advice, Sanna Koskiniemi for advice on library construction, and Irmeli Barkefors for assistance with figure creation and careful proof-reading of the manuscript.

Jean-Marc Ghigo provided us with the strain MFD*pir*. pMarC9-R6k was a gift from George Church (Addgene plasmid #89477; http://n2t.net/addgene:8947; RRID:Addgene_89477). Transposon sequencing of the whole libraries was performed by the SNP&SEQ Technology Platform in Uppsala. The facility is part of the National Genomics Infrastructure (NGI) Sweden and Science for Life Laboratory. The SNP&SEQ Platform is also supported by the Swedish Research Council and the Knut and Alice Wallenberg Foundation. This study was made possible by grants from the ERC (Advanced grant no. 885360); the Swedish Research Council (grant no. 2016-06213, 2018-03958, and 2023-03442); the Knut and Alice Wallenberg Foundation (grant no. 2016.0077, 2017.0291 and 2019.0439); and the eSSENCE e-science initiative. Some of the computations and data management were enabled by resources provided by the Swedish National Infrastructure for Computing (SNIC) at UPPMAX, partially funded by the Swedish Research Council through grant agreement no. 2022-06725.

## Author contributions

O.B.: strain construction, sequencing, input on microfluidic device design, microfluidics experiments, single-cell isolation, analysis, code development, writing. P.K.: device design, code development, analysis, single-cell isolation, writing. E.A.: built the optical setup including the optical tweezer, custom pneumatics, input on device design. M.T. was involved in the early phase of the device design. J.E.: conceived the idea of single-cell isolation based on a transposon library, input on device design, supervision, writing, and interpretation.

## Code and data availability

All data, analysis output, and code will be available on the SciLifeLab data repository, BioImage Archive, European Nucleotide Archive, and GitHub upon publication or upon reasonable request. Strains are available upon reasonable request.

## Extended Data

**Extended Data Fig. 1.**
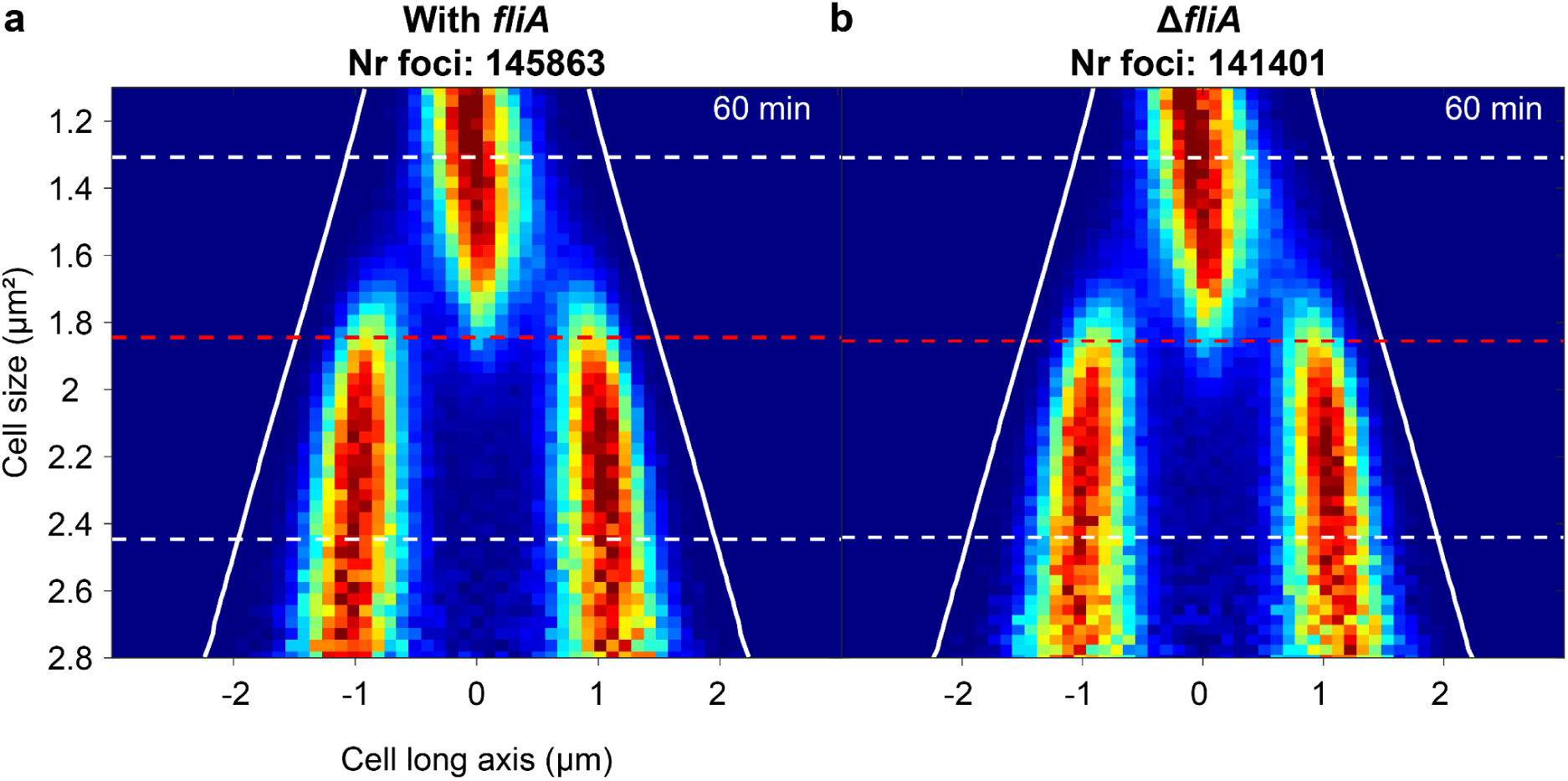
Effect of *fliA* deletion on replication initiation control. **a**, Fork plot of a strain with *fliA*. **b,** Fork plot of a strain without *fliA*. The white dashed lines correspond to the average birth or division area. Lines are otherwise the same as in Fig. 1d. The average generation time is highlighted in the top-right corner of each fork plot.

**Extended Data Fig. 2.**
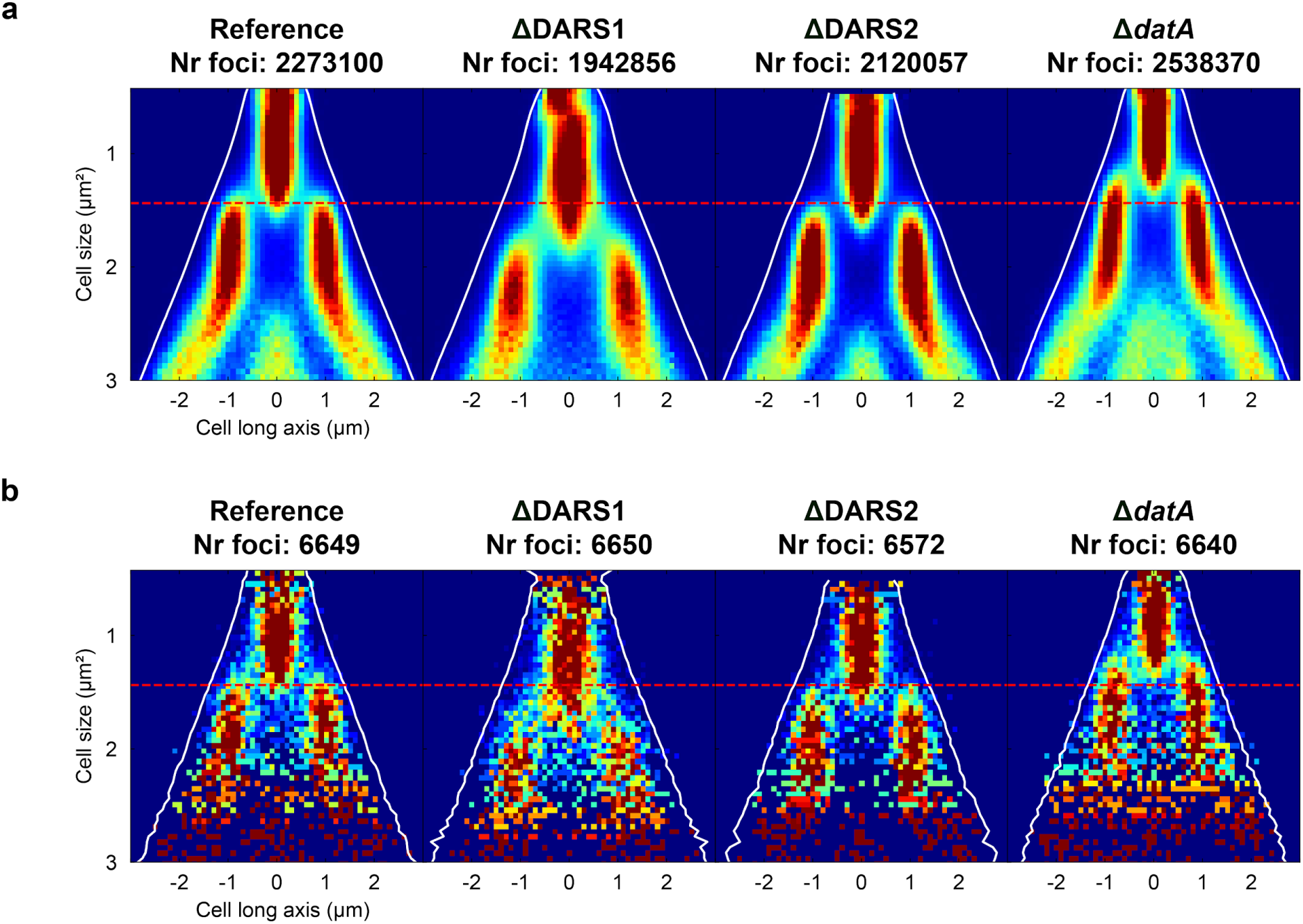
Full fork plots of the arrayed DnaA-ATP/ADP regulatory knockouts. **a**, All-traps fork plots. **b,** Downsampled fork plots based on **(a)**. Lines as in Fig. 2a.

**Extended Data Fig. 3.**
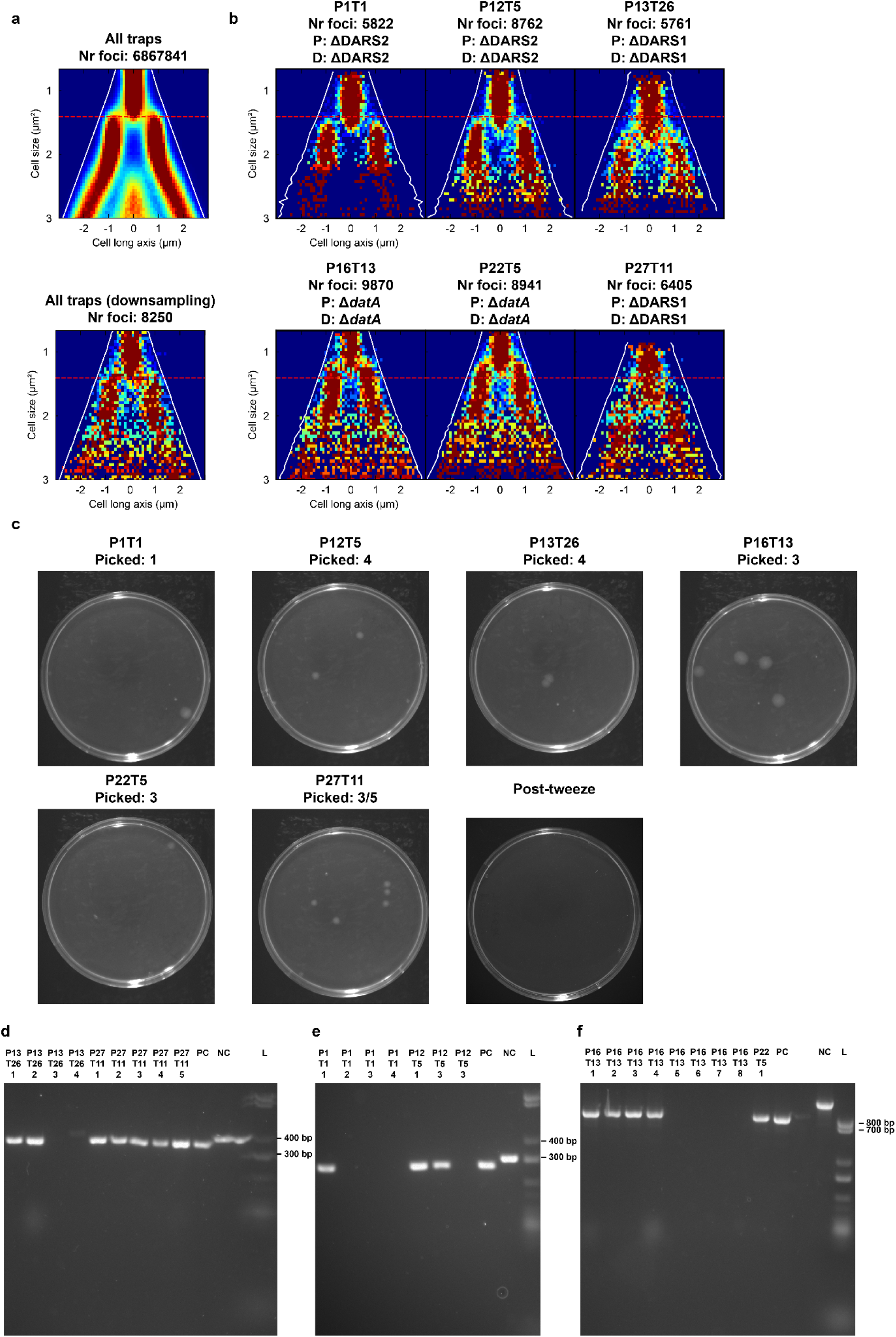
Pooled DnaA-ATP/ADP regulatory knockouts single-cell isolation. **a**, Top: Fork plots for all traps. Bottom: Downsampled fork plot based on the all traps one. **b,** Single-trap fork plots where cells were isolated from. **c,** Agar plates where cells were collected. “Picked” indicates the number of cells isolated from a particular trap. “/” indicates ambiguity in the number picked. **d–f,** Agarose gels with PCR products from each set of control primers for the different mutants. PC: positive control with the deletion. NC: negative control without the deletion. L: DNA ladder. All wells without product are likely not *E. coli* colonies. **d,** ΔDARS1 gel. **e,** ΔDARS2 gel. **f,** Δ*datA* gel. Labels and lines as in Fig. 3a–c.

**Extended Data Fig. 4.**
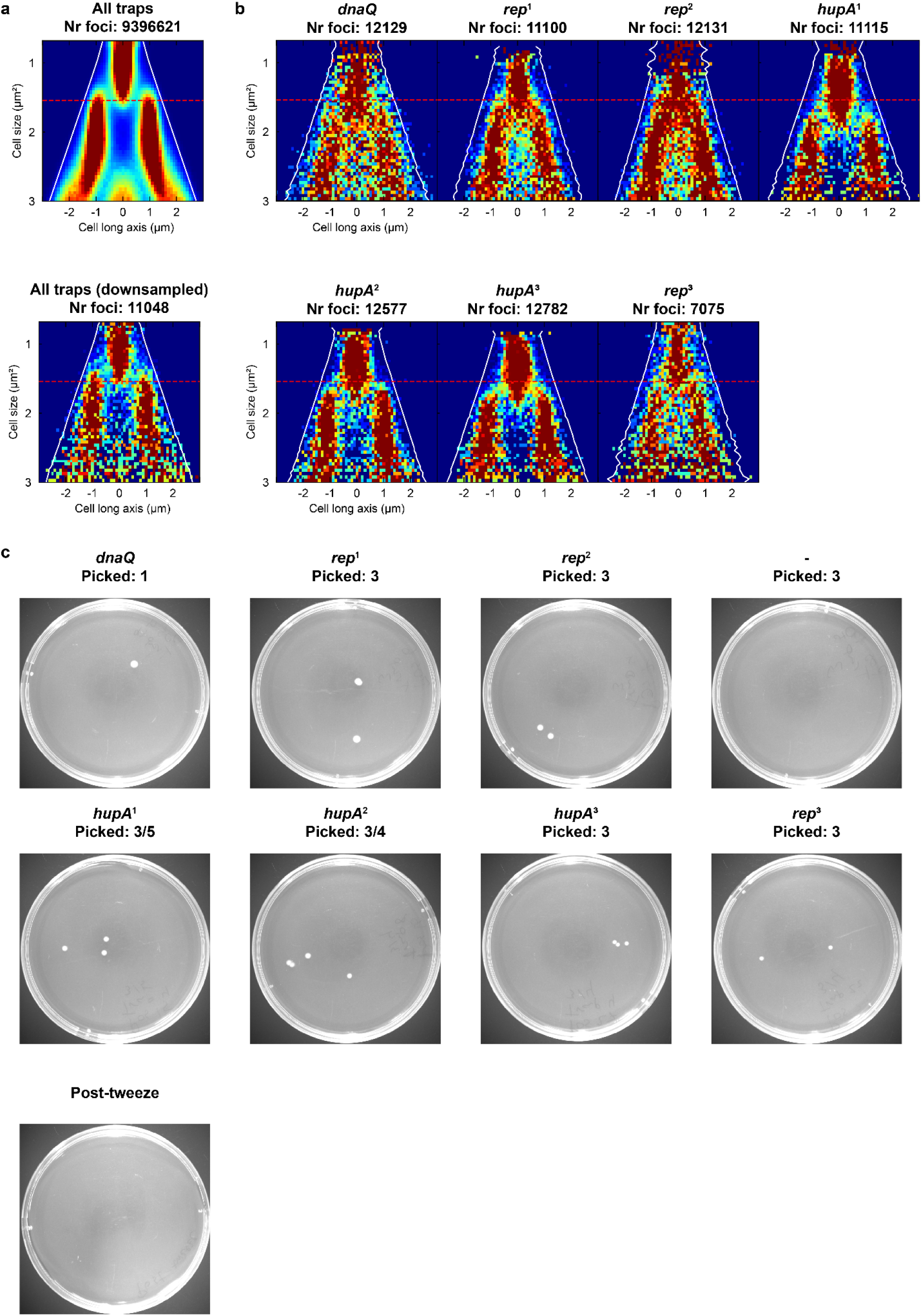
Pooled CRISPRi library single-cell isolation. **a**, Top: Fork plot for all traps. Bottom: Downsampled fork plot based on **(a)**. **b,** Single-trap fork plots where cells were isolated from. **c,** Agar plates where cells were collected. Lines and labels as in Figs. 3d–f and Extended Data Fig. 3c.

**Extended Data Fig. 5.**
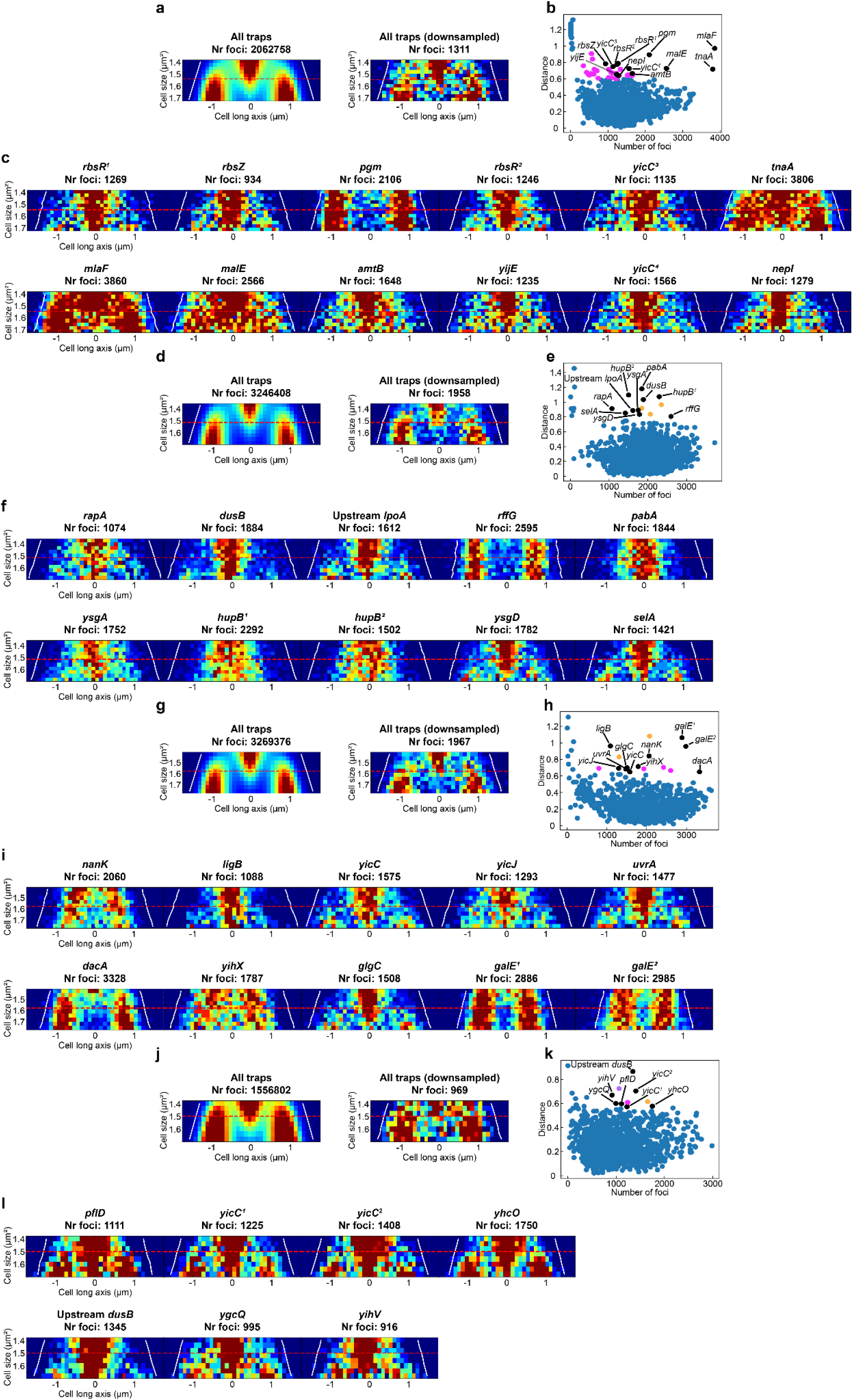
Isolation of single cells from the transposon mutant library replicates not shown in. Fig. 4**. a, d, g, j,** Left: Initiation fork plot for all traps. Right: Downsampled initiation fork plot from the all-traps fork plot. **b, e, h, k,** Same structure as Fig. 4b. **c, f, i, l,** Initiation fork plots for the traps with successful cell isolation. Footnotes indicate mutants that appear in the same replicate at least twice. **a–c,** Replicate 2. The transposon sites are not the same for the *rbsR* and *yicC* mutants, respectively. **d–f,** Replicate 3. The transposon sites are not the same for the *hupB* mutants. **g–i,** Replicate 4. The two *galE* mutants have the same transposon insertion site. **j–l,** Replicate 5. The transposon sites are not the same for the *yicC* mutants.

**Extended Data Fig. 6.**
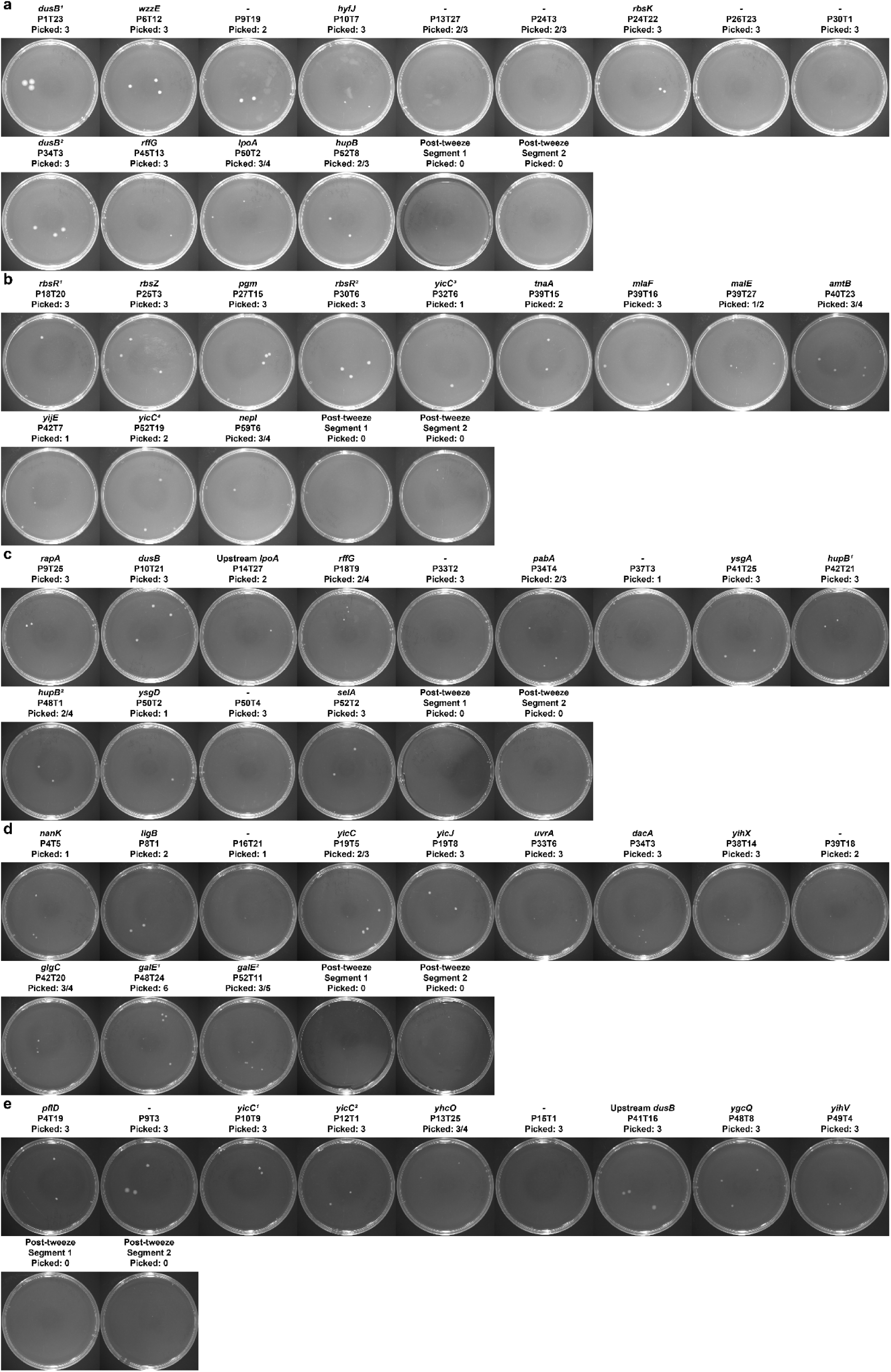
Agar plate for collection of isolated cells from transposon library screens. **a**, Replicate 1 (Fig. 4). **b,** Replicate 2. **c,** Replicate 3. **d,** Replicate 4. **e,** Replicate 5. Plates with unsuccessful isolation are included, indicated by -. Labels as in Extended Data Fig. 3c.

**Extended Data Fig. 7.**
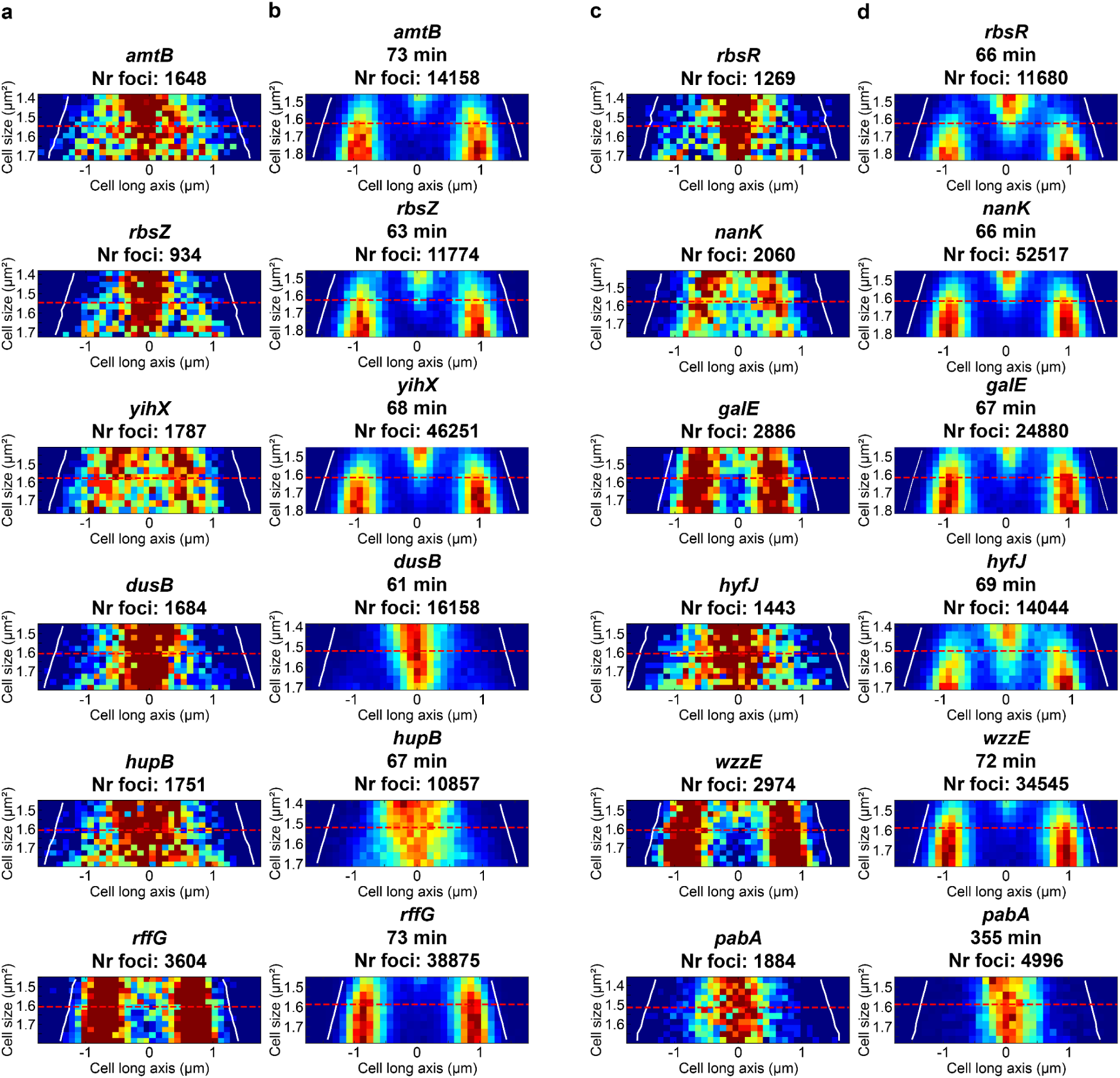
All pooled and arrayed single-mutant comparisons. a and c, Single-trap initiation fork plots. **b and d,** Single-trap initiation fork plots for multiple traps. Lines as in Fig. 5.

