## Supplementary material for "Optical pooled screening of a bacterial transposon mutagenesis library": SI

#### Supplementary Note 1: Increasing throughput of the pooled screen

We have used at most two microfluidic segments, including 1,680 traps, out of the 14 segments, including 11,760 traps, for our pooled screens. All segments can be used if we connect the fluidic ports of each segment in an additional layer to avoid an unmanageable number of interfaces to separate media sources. However, if all segments are utilized, the screens would have to be longer to allow the analysis of our particular phenotype. This is because we need to accumulate enough data per trap to robustly call deviating phenotypes while being limited to how quickly images can be acquired, how fast the microscope stage can move, and the speed of the real-time analysis. Using an objective with lower magnification would reduce the number of fields of view that have to be imaged, but the magnification cannot be too low, as we still need enough spatial resolution to create fork plots.

#### Supplementary Note 2: Repeat of arrayed DnaA-ATP/ADP regulatory knockout screen

Compared with the replicate shown in Fig. 2, the replicate in Supplementary Fig. 2 had more traps from the knockouts within the convex hull enveloping the reference strain traps, fewer detected foci, and more variability in the distance for the reference strain and the DARS1 knockout. This is due to the fact that the two mentioned strains started growing slower and more or less stopped replicating for approximately half of the screen (source data available on request). Why this happened we are not sure, but it only affected these two strains as they are on a separate microfluidic device from the DARS2 and *datA* knockouts. This second device did not have the same issues. Due to this problem with the reference strain, we applied the 300 foci threshold for the number of detected foci when constructing the convex hull, resulting in less overlap with knockout traps. Applying the threshold for the replicate in Fig. 2 is not necessary, as all of the reference strain traps have more than 300 detected foci.

### Supplementary Note 3: Small-scale library cell isolation replicates

In the replicate for the pooled DnaA-ATP/ADP regulatory mutants, we isolated cells from eight different traps, again to cover all mutants but now also the reference strain. The pooled fork plot for all traps appeared as the reference strain (Supplementary Fig. 3a and 3b). The traps where cells were isolated from were chosen based on their distance from the cluster center (Supplementary Fig. 3c), with the predicted reference strain traps' distance being low and their visual similarity to Fig. 2a. Our predicted genotypes mostly agreed with the determined genotypes (Supplementary Fig. 3d and 3e). For P18T15, we predicted the genotype to be Δ*datA*, while the determined genotype was no deletions. As P18T15's phenotype is similar to P16T26, we assume that we accidentally picked cells from the wrong trap or that the trap somehow had been repopulated with the reference strain in the time between image acquisition finishing and single-cell isolation (P18T15 was the fifth trap done in the day). Furthermore, there were 9 colonies on the plate, while we only isolated 3 cells. Where the extra cells came from we do not know. Agarose gels for the genotyping can be seen in Supplementary Fig. 3g–k.

In the replicate of the CRISPRi library screen, the pooled fork plots for all of the traps were reference-like (Supplementary Figs. 4a and 4b). Out of the seven isolated knockdowns (Supplementary Fig. 4c–f), most were classified by Camsund et al.[1] as having an effect on initiation control or replication as a whole (*hupA*, *diaA*, *rep*, and *dnaQ*). However, we also isolated the *mrr* knockdown and the reference strain. Camsund et al.[1] did not classify the *mrr* knockdown as having a smaller initiation size (Supplementary Figs. 4d and 4e). It could be that the trap in our screen just happens to display a different phenotype and that potential other *mrr* traps appear reference-like. Another trap produced an ambiguous result, with some colonies being classified as *diaA* and others as reference (ref) (Supplementary Fig. 4d and 4e). The fork plot from this trap looks similar to the other *diaA* trap, so the correct genotype is likely *diaA* and a few reference cells we collected by mistake.

### Supplementary Note 4: Unsuccessful single-cell isolation and transposon mapping

We were unable to recover any cells from 12 out of 93 traps (Extended Data Fig. 4c, Extended Data Fig. 6, and Supplementary Fig. 1d). We can only speculate why recovery was unsuccessful, but it could be that the cells got stuck in the connector or tubing during ejection of liquid onto the agar plate or that cells were damaged by the IR laser used as the optical tweezer. For the transposon mutagenesis pooled screens in Fig. 4 and Extended Data Fig. 5d–l, there were 6 traps that were classified as outliers, but we never ended up isolating cells from them. This was because there was an accumulation of cells surrounding those traps that, for unknown reasons, were stuck to either the PDMS or coverslip. These cells could not be moved by the tweezer or flushed away by increasing the flow. Accessing these traps therefore became impossible, hence why we did not isolate cells from them. In the screen in Extended Data Fig. 5a–c, too many traps (36) were within the threshold to feasibly isolate cells from in a day. Traps were chosen based on their phenotype so that, in total, larger, smaller, and variability in initiation size were covered.

In the transposon mutagenesis pooled screens, there were 2/59 traps from which we successfully recovered cells, but we did not do transposon mapping. This is because there is a ~20% risk that the conjugative suicide vector gets integrated into the chromosome when using conjugation to deliver transposons[2]. Mutants with vector integration express the transposase constitutively, meaning that we cannot link a phenotype to the disruption of a particular genomic region. Since this is an issue for us, unlike traditional TIS where these mutants can be filtered out during analysis[2], we introduced a *catsacB* cassette into our vector. The cassette consists of the *cat* gene, which confers chloramphenicol resistance, and the *Bacillus subtilis* gene *sacB*, which confers sucrose sensitivity. By counterselecting on sucrose, we could reduce the frequency of exconjugants with whole-plasmid integration. However, the counterselection is not 100% efficient, so our library contains a fraction of mutants carrying the transposase. To avoid spending time mapping transposons for these mutants, we checked for the presence of the MarC9 gene by PCR, and if it was present, we did not proceed with transposon mapping.

#### Supplementary Note 5: Per-trap distance analysis of the individual transposon mutants

Besides the *hyfJ* mutant, there were mutants with respective insertions in *amtB*, *rbsZ*, *yihX*, *rbsR*, *nanK*, and *galE* that had inconsistent results between the pooled and arrayed screens. One possible reason for this is that there is phenotypic heterogeneity within each mutant, which can be seen for the *galE* mutant (Supplementary Fig. 7). To investigate this, we performed per-trap distance analysis on the arrayed screens of these mutants (Supplementary Fig. 6). We also included the *dusB*, *hupB*, *rffG*, *wzzE*, and *pabA* mutants to see if there was heterogeneity for them as well. The analysis was done the same way as for the arrayed DnaA-ATP/ADP regulatory mutants. What we are looking for is if the mutants are farther away from the reference cluster center compared to most reference traps and how much this distance varies between traps from the same mutant.

The *pabA*, *dusB*, and *hupB* mutants all separate well from the reference strain in their respective screen, which is to be expected given that their phenotypes are significantly different from the reference strain. The *rffG* mutant separated from the reference cluster, but, in absolute numbers, not as much as in the pooled screen (compare Supplementary Fig. 6c and Fig. 4b). This could stem from the phenotypic difference being smaller in the arrayed screen and there being less data in the arrayed screen compared to the pooled screen. The *wzzE* mutant, on average, barely separates from the reference, likely because its phenotype is less pronounced in relation to the reference strain compared to the *rffG* mutant (Fig. 5, Extended Data Fig. 7, and Supplementary Fig. 6c). The *amtB*, *rbsZ*, *nanK*, and *yihX* mutants look similar to the reference strain in their respective experiments (Supplementary Fig. 6a and 2d), which, given their function, makes sense. The *rbsR*, *hyfJ*, and *galE* mutants are more interesting due to more heterogeneity between traps (Supplementary Figs. 6a, 6b, and 6d). Given the fork plot for all traps for the *rbsR* mutant is similar to the reference strain (Extended Data Fig. 7d and Supplementary Fig. 5a), it is surprising to see such heterogeneity between traps. One explanation for this could be that the screen duration was not as long as the other arrayed screens (4 h instead of 6 or 7 h), resulting in there not being enough data per trap to get a representative fork plot. If, on the other hand, there is heterogeneity in the mutant, the trap in the pooled screen could appear similar to the reference strain by chance. For the *hyfJ* mutant, its fork plot appears slightly blurrier than the corresponding reference strain (Supplementary Fig. 5b), which could explain the heterogeneity that can be seen. Some of the traps that are further away from the reference cluster could have a larger initiation size, similarly to the *hyfJ* trap in the pooled experiment. For the *galE* mutant, most traps have a distance similar to individual reference traps (Supplementary Fig. 6d). The few whose distance is further away have few foci within the size range used for the initiation fork plots. However, due to the heterogeneity explained in the section *"Discussion of individual transposon mutants"*, some cells are smaller and others larger. This results in data from these cells being outside of the reference initiation size range, so these traps appear to have little data in our analysis. Whether there is any underlying biology that can explain the heterogeneity between traps for these three mutants, and if so what that biology looks like, requires further investigation to elucidate.

### Supplementary Note 6: Discussion of individual transposon mutants

When we performed arrayed screens of the different transposon mutants (Extended Data Fig. 7 and Supplementary Fig. 5) and compared the results to the pooled screens, we observed that some mutants had similar phenotypes (*dusB*, *hupB*, *rffG*, *wzzE*, and *pabA*), while others were dissimilar (*amtB*, *rbsZ*, *yihX*, *rbsR*, *nanK*, *hyfJ*, and *galE*). Here follows a discussion on the different mutants.

Previously, a dCas9 knockdown of *hupB* has been shown to result in a larger initiation size[1], which is consistent with our results. It is therefore reasonable that the results are consistent across the pooled and arrayed screens. For the *dusB* mutant, we assume that, since *dusB* is in the same operon as *fis*, the expression of *fis* is also altered, resulting in the observed phenotype. This is despite our *dusB* mutant not carrying transposons in the RNA sequence elements that enable higher translation of *fis* compared to *dusB*[3]. The phenotype of our *dusB* mutant also resembles a knockdown of *fis*[1], in support of our assumption. That both the *hupB* and *dusB* mutants are consistent with one another between pooled and arrayed screens shows that the pooled screens can capture known relationships between phenotype and genotype.

For *pabA*, there are no prior measurements regarding replication, but the gene can be indirectly linked to replication. PabA is involved in the synthesis of 4-amino-4-deoxychorismate (ADC). ADC is a precursor of p-aminobenzoate (PABA), which later is processed into folate[4]. Folate is needed for nucleotide synthesis. Our medium does not contain folate, meaning that the mutant likely cannot synthesize enough nucleotides to maintain normal replication speed. This explains the fork plot phenotype (Fig. 5, Extended Data Fig. 7, and Supplementary Fig. 5c) and the long generation time (355 min) (Extended Data Fig. 7 and Supplementary Fig. 5c). For *rffG* and *wzzE*, there are no direct or indirect connections between their function and replication. Interestingly, the *rffG* and *wzzE* mutants have similar phenotypes (Figs. 4 and 5), which may be because they are in the same operon[5] and are both involved in enterobacterial common antigen synthesis[6]. The phenotype (Figs. 4 and 5, Extended Data Fig. 7, and Supplementary Fig. 1C), at least for the *rffG* mutant, stems from the transposon insertion rather than any other mutation (confirmed by whole-genome sequencing; the *wzzE* mutant has not been whole-genome sequenced).

However, the generation times of the two mutants are longer than the reference strain (65 min for the reference, and 72 and 73 min for the *wzzE* and *rffG* mutants, respectively) (Extended Data Fig. 7 and Supplementary Fig. 5c), so whether the initiation phenotype we observe is due to this discrepancy or not requires further studies.

The mutants with inconsistent results between the pooled and arrayed screens (*amtB*, *rbsZ*, *yihX*, *rbsR*, *nanK*, *hyfJ*, and *galE)* have no obvious connection to initiation control. *amtB* is involved in ammonium transport during nitrogen starvation[7]. Our cells should not be nitrogen-starved, as our minimal medium is supplemented with amino acids. A nonsense AmtB protein should therefore not have much of an effect on our cells, especially in terms of replication. The phenotype observed in the pooled screen is unexpected, and we cannot explain it. *rbsR* and *rbsZ* are both located in the *rbs* operon, which is involved in ribose catabolism and transport. RbsZ is a regulatory RNA that interacts with another regulatory RNA (RybB). These are, however, more abundant in stationary phase[8]. It is not clear how their interplay would affect initiation control. RbsR is the transcription fator for the *rbs* operon and negatively regulates itself. It also regulates the *purHD* operon through repression, and *add* and *udk* through activation[9]. The former is involved in purine biosynthesis, and the latter is involved in purine salvage and degradation. Given that ribose is central to nucleotide biosynthesis, the observed effect could be due to altered replication speed.

The *yihX*, *nanK*, and *hyfJ* mutants were all whole-genome sequenced, and their pooled and arrayed screens had inconsistent results. The *nanK* and *yihX* transposon mutants had not acquired any additional mutations. These two genes are involved in different metabolic processes (*nanK* in N-acetylmannosamine metabolism and *yihX* in α-D-glucose-1-phosphate metabolism)[10,11]. Any possible connection with initiation control would be indirect through various metabolic pathways. *hyfJ* is a part of the *hyf* operon, an operon coding for different subunits in a hydrogenase. The operon is induced under fermentative conditions by formate[12]. Our growth condition is not fermentative, so we did not expect to observe any phenotype due to the transposon insertion, especially on initiation control. The phenotype in the pooled screen was therefore unexpected. However, in the arrayed screen, the phenotype differed not only from the pooled screen (Fig. 5) but also from the reference strain in the arrayed screen (Supplementary Fig. 5b), which was unexpected. This can perhaps be attributed to a point mutation in *pntA*, which resulted in the amino acid substitution D314V. *pntA* codes for the α subunit in pyridine nucleotide transhydrogenase, which is involved in NADPH synthesis[13]. Whether this mutation causes loss of function is not known. Moreover, there is no known connection between *hyfJ* and *pntA*, and *pntA* itself is not known to be involved in initiation control. It remains to be seen if there is any connection between these two genes in regard to initiation control or any other biological process.

The last mutant has its transposon in *galE*. *galE* is part of the *gal* operon, which ensures galactose metabolism[14]. Since our medium does not contain galactose, we did not expect to observe any effect on initiation control or growth when disrupting *galE*. However, in the pooled screen, we observed that the mutant with a transposon insertion in *galE* exhibited distinct morphology and appeared darker in phase-contrast (Supplementary Fig. 7a). We assume that the initiation control phenotype is related to these differences. Interestingly, in the arrayed screen, we observed heterogeneity among traps, with cells in some traps resembling the pooled screen phenotype while others appeared normal (Supplementary Fig. 7b). The cause of this heterogeneity warrants further studies.

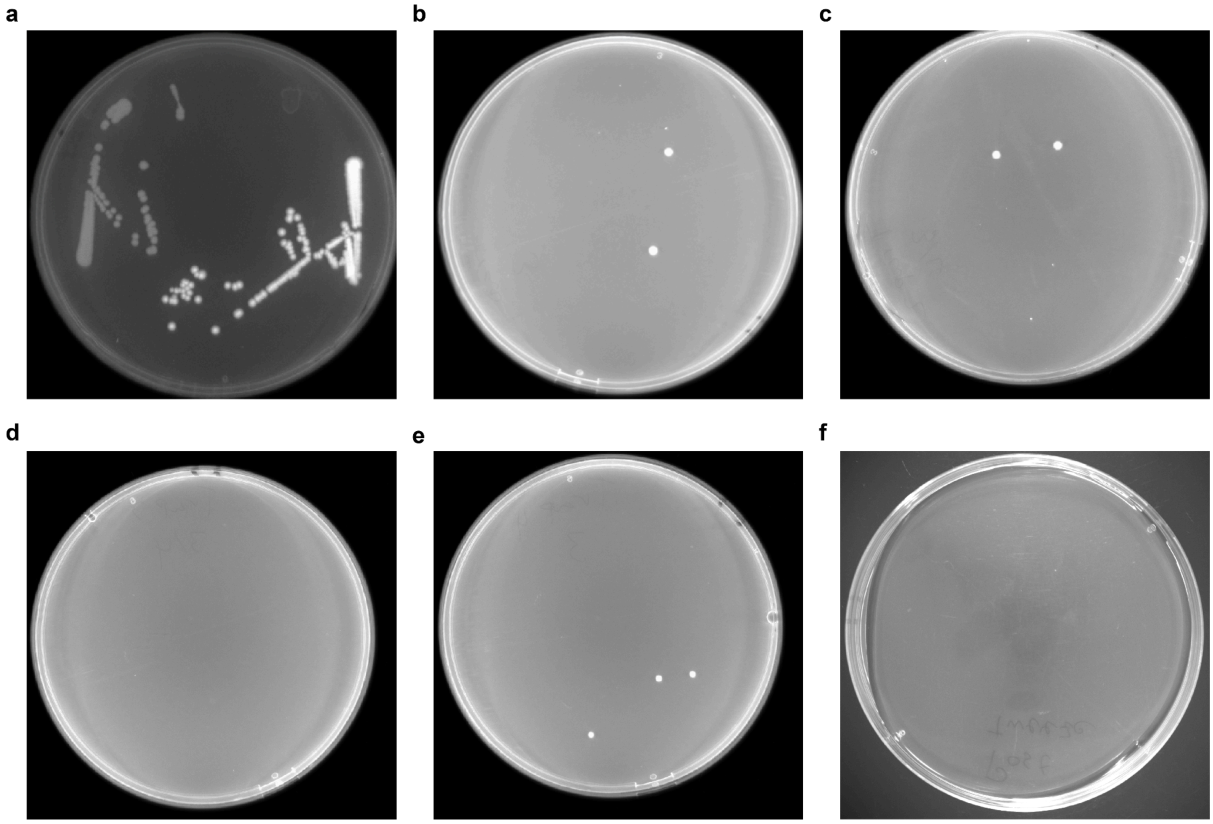


**Supplementary Fig. 1 | Isolation of fluorescent cells from a mixed population. a,** Non-fluorescent and fluorescent cells streaked from glycerol stock onto an LA plate. **b,** LA plate for trap 1. **c,** LA plate for trap 2. **d,** LA plate for trap 3. **e,** LA plate for trap 4. **f**, LA plate after the isolations shown in **(b–e)**. Medium was collected overnight.

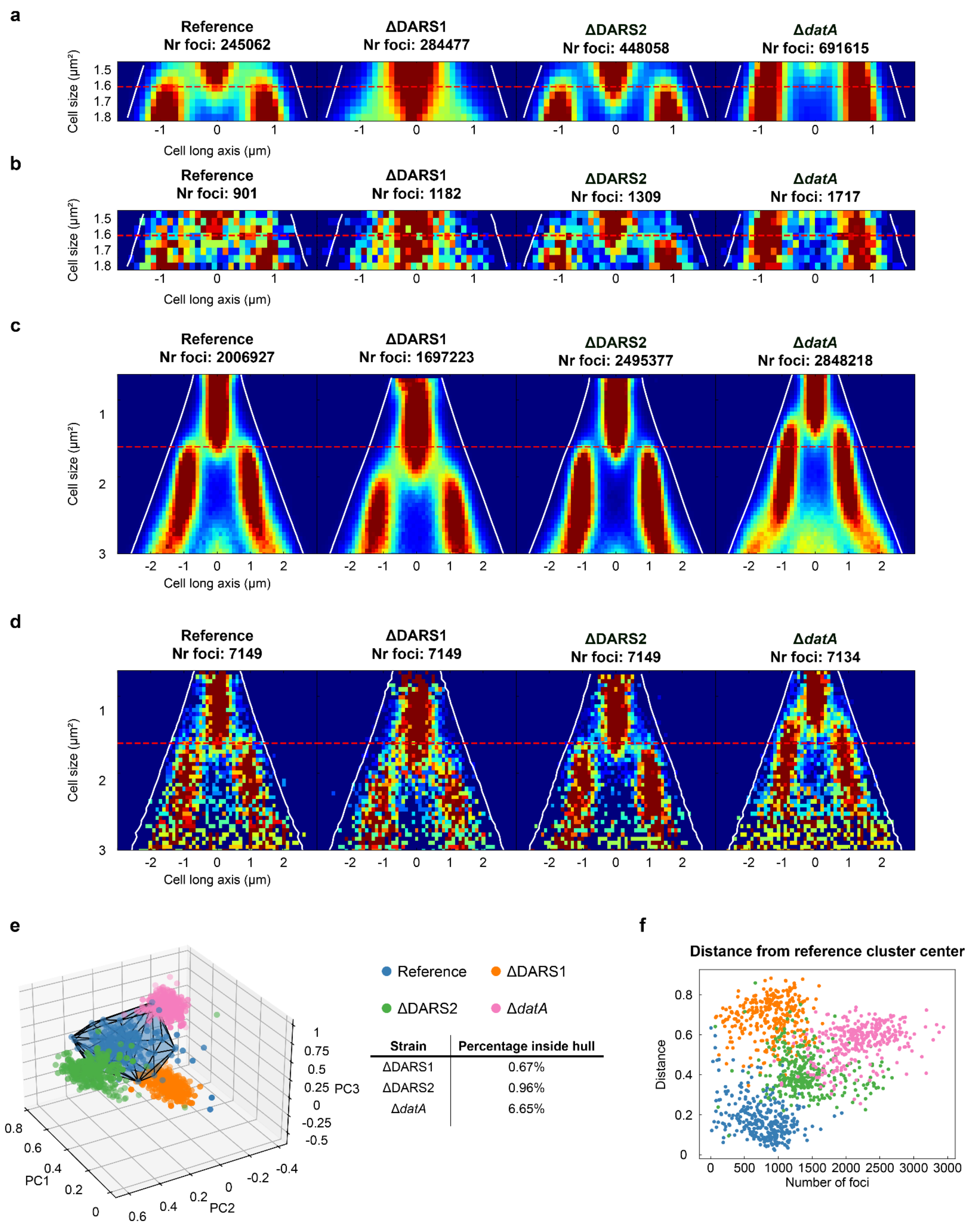


**Supplementary Fig. 2 | Replicate of arrayed DnaA-ATP/ADP regulatory mutant screen. a,** Initiation fork plots for all traps. **b,** Downsampled initiation fork plots based on **(a)**. **c,** Fork plot for all traps. **d,** Downsampled fork plots based on **(c)**. **e,** PCA of the initiation fork plots for each trap. A convex hull surrounds the reference strain traps (blue) that had more than 300 foci. The table indicates the percentage of traps of the DnaA-ATP/ADP knockouts (ΔDARS1: orange; ΔDARS2: green; Δ*datA*: pink) that fall within the convex hull. **f,** Euclidean distance for each trap from the center of the principal components of all reference traps.

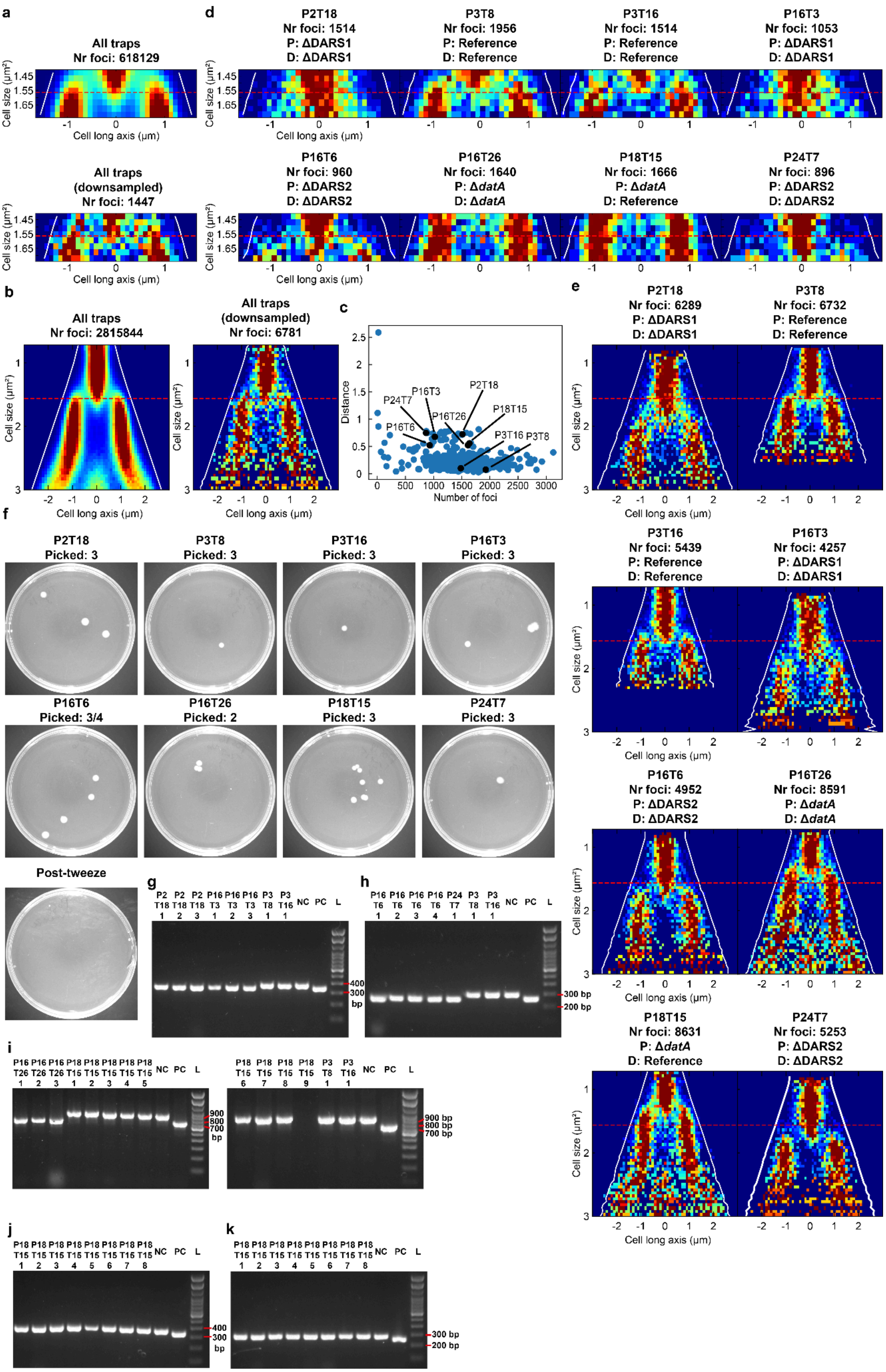
a
All traps
Nr foci: 618129
Cell size (µm²)
Cell long axis (µm)
All traps
(downsampled)
Nr foci: 1447
d
P2T18
Nr foci: 1514
P: ΔDARS1
D: ΔDARS1
P3T8
Nr foci: 1956
P: Reference
D: Reference
P3T16
Nr foci: 1514
P: Reference
D: Reference
P16T3
Nr foci: 1053
P: ΔDARS1
D: ΔDARS1
P16T6
Nr foci: 960
P: ΔDARS2
D: ΔDARS2
P16T26
Nr foci: 1640
P: ΔdatA
D: ΔdatA
P18T15
Nr foci: 1666
P: ΔdatA
D: Reference
P24T7
Nr foci: 896
P: ΔDARS2
D: ΔDARS2
b
All traps
Nr foci: 2815844
All traps
(downsampled)
Nr foci: 6781
c
Distance
Number of foci
P24T7
P16T3
P16T26
P2T18
P18T15
P16T6
P3T16
P3T8
e
P2T18
Nr foci: 6289
P: ΔDARS1
D: ΔDARS1
P3T8
Nr foci: 6732
P: Reference
D: Reference
P3T16
Nr foci: 5439
P: Reference
D: Reference
P16T3
Nr foci: 4257
P: ΔDARS1
D: ΔDARS1
P16T6
Nr foci: 4952
P: ΔDARS2
D: ΔDARS2
P16T26
Nr foci: 8591
P: ΔdatA
D: ΔdatA
P18T15
Nr foci: 8631
P: ΔdatA
D: Reference
P24T7
Nr foci: 5253
P: ΔDARS2
D: ΔDARS2
f
P2T18
Picked: 3
P3T8
Picked: 3
P3T16
Picked: 3
P16T3
Picked: 3
P16T6
Picked: 3/4
P16T26
Picked: 2
P18T15
Picked: 3
P24T7
Picked: 3
Post-tweeze
g
NC PC L
400
300
bp
h
300 bp
200 bp
i
900
800
700
bp
900 bp
800 bp
700 bp
j
k
400
300
bp
300 bp
200 bp

**Supplementary Fig. 3 | Replicate of DnaA-ATP/ADP regulatory mutants pooled screen. a,** Top: Initiation fork plot for all traps. Bottom: Downsampled initiation fork plot. **b,** Left: Full fork plot for all traps. Right: Downsampled fork plot for all traps. **c,** Euclidean distance from the PCA cluster center for all traps. Black points indicate the traps from which cells were isolated. **d,** Initiation fork plots of the traps where cells were isolated from. The name of each trap and the number of foci are indicated above each plot. The predicted genotype is indicated with P, and the determined genotype by colony PCR is indicated with D. **e,** Full fork plots for the traps where cells were isolated from. Same notation as in **(d)**. **f,** Agar plates where cells were collected. "Picked" indicates the number of cells isolated from a particular trap. "/" indicates ambiguity in the number of picked cells. **g–k,** Agarose gels with PCR products from each set of control primers for the different mutants. PC: positive control with the deletion. NC: negative control without the deletion. L: DNA ladder. **g,** ΔDARS1 gel. **h,** ΔDARS2 gel. **i,** Δ*datA* gel. P18T15 9 is likely not an *E. coli* colony; hence there is no PCR product. **j,** Gel to check whether P18T15 is ΔDARS1. **k,** Gel to check whether P18T15 is ΔDARS2. **j and k**, P18T15 9 was not included (see reason in **(i)**). Labels and lines as in Fig. 3a–c.

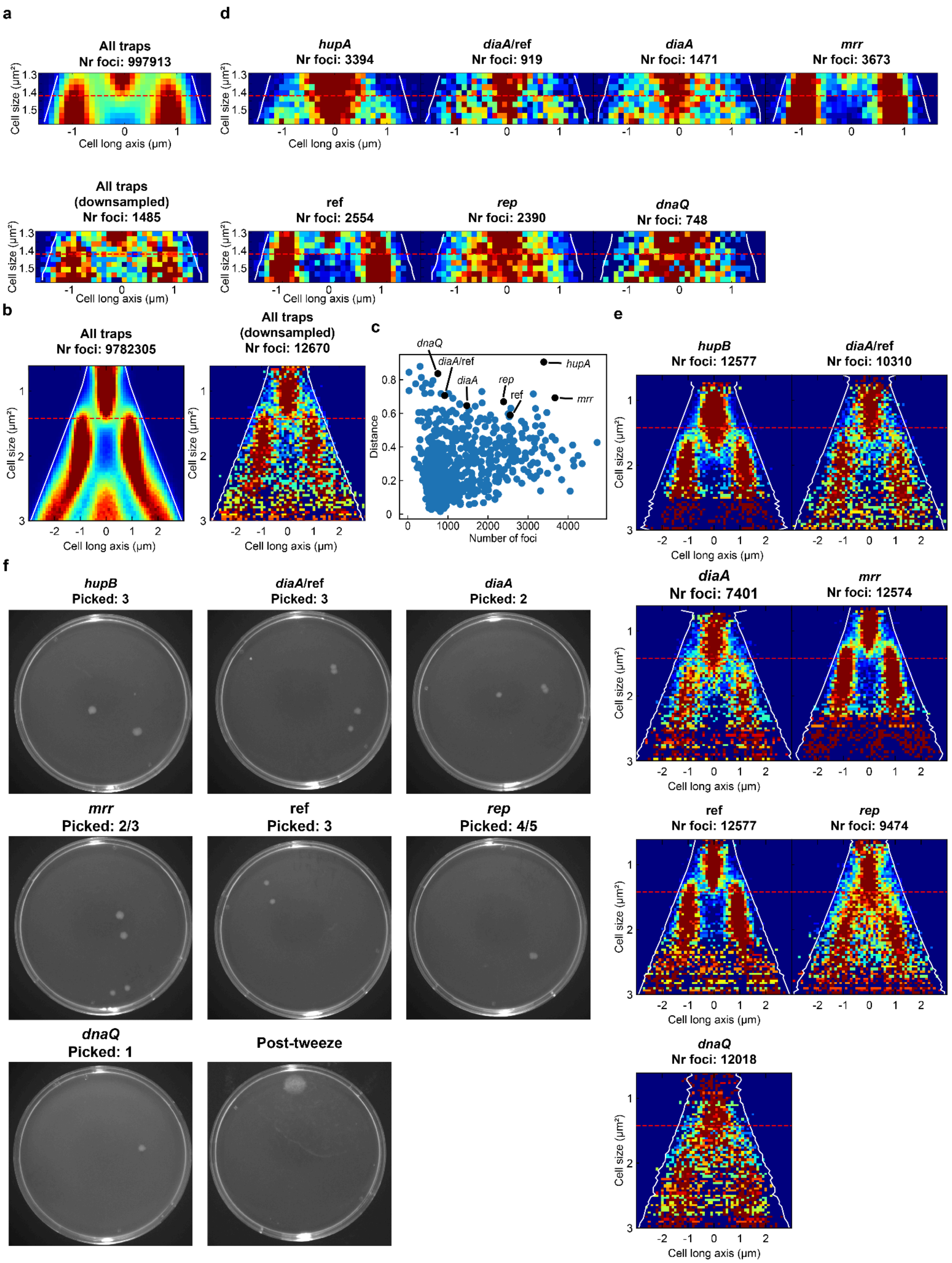


**Supplementary Fig. 4 | Replicate of CRISPRi library pooled screen. a,** Top: Initiation fork plot of all traps. Bottom: Downsampled initiation fork plot. **b,** Left: Full fork plot of all traps. Right: Downsampled full fork plot. **c,** Euclidean distance from the PCA cluster center for all traps. Black points indicate the traps from which cells were isolated. **d,** Initiation fork plots of the traps where cells were isolated from. The determined sgRNA target and the number of detected foci are indicated above each plot. **e,** Full fork plots for the traps where cells were isolated from. Same notation as in **(c)**. **f,** Agar plates where cells were collected. “Picked” indicates the number of cells isolated from a particular trap. “/” indicates ambiguity in the number of picked cells.

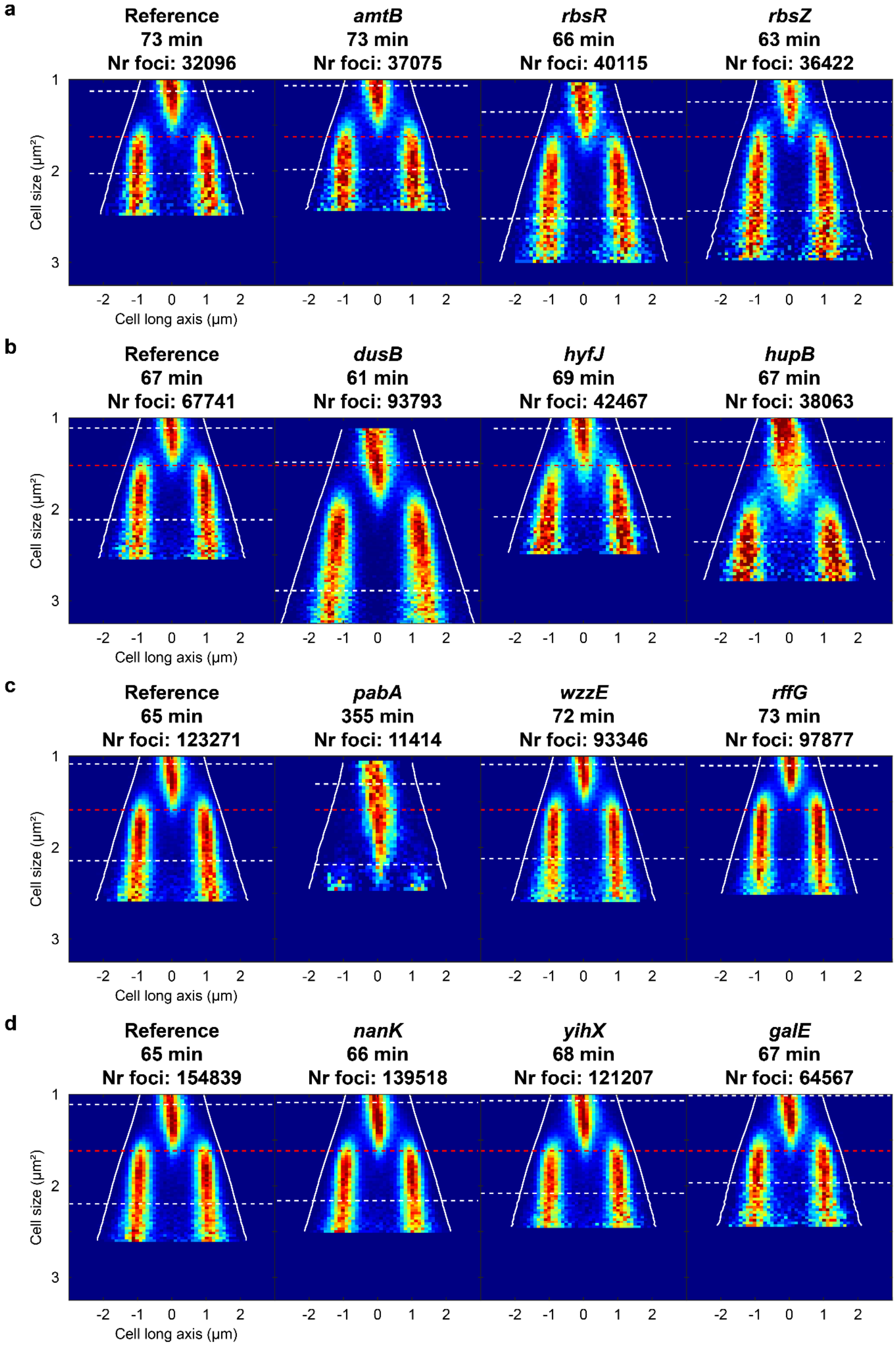

a
Reference
73 min
Nr foci: 32096
amtB
73 min
Nr foci: 37075
rbsR
66 min
Nr foci: 40115
rbsZ
63 min
Nr foci: 36422
Cell size (µm²)
Cell long axis (µm)
b
Reference
67 min
Nr foci: 67741
dusB
61 min
Nr foci: 93793
hyfJ
69 min
Nr foci: 42467
hupB
67 min
Nr foci: 38063
c
Reference
65 min
Nr foci: 123271
pabA
355 min
Nr foci: 11414
wzzE
72 min
Nr foci: 93346
rffG
73 min
Nr foci: 97877
d
Reference
65 min
Nr foci: 154839
nanK
66 min
Nr foci: 139518
yihX
68 min
Nr foci: 121207
galE
67 min
Nr foci: 64567

**Supplementary Fig. 5 | Full fork plots of transposon mutants and the reference strain grown separately. a–d,** Fork plots of all transposon mutants imaged separately and the reference strain imaged in the same experiment. The red dashed line corresponds to the average initiation size of the reference strain in the same experiment. **a,** Arrayed screen 1. **b,** Arrayed screen 2. **c,** Arrayed screen 3. **d,** Arrayed screen 4.

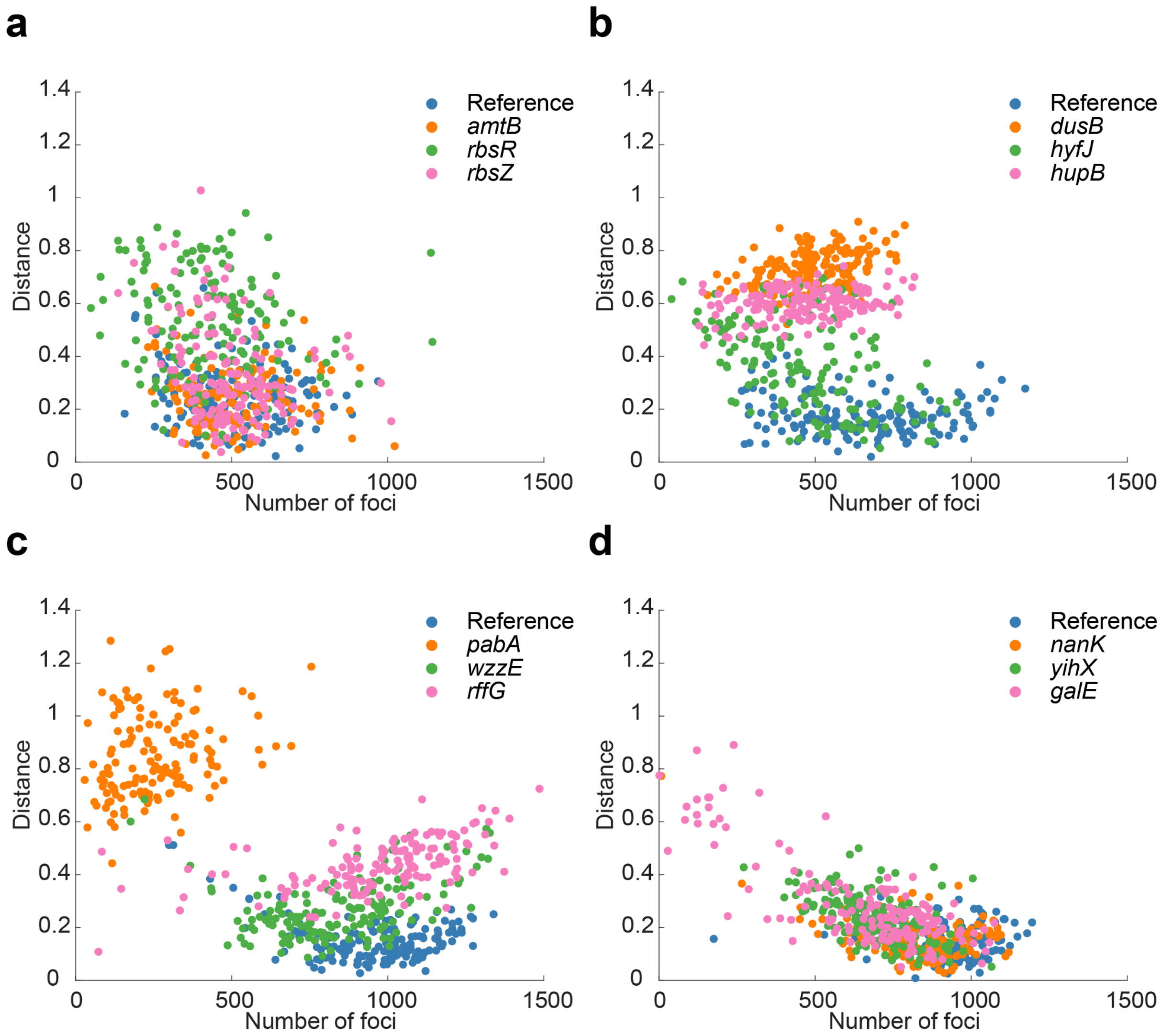


**Supplementary Fig. 6 | Per-trap distance analysis on arrayed transposon mutants.** Euclidean distance for each trap from the center of the principal components of all reference traps. Four separate arrayed screens with mutants found in the screens shown in Extended Data Fig. 7 and Supplementary Fig. 5. Data comes from cells that have not been tracked.

a

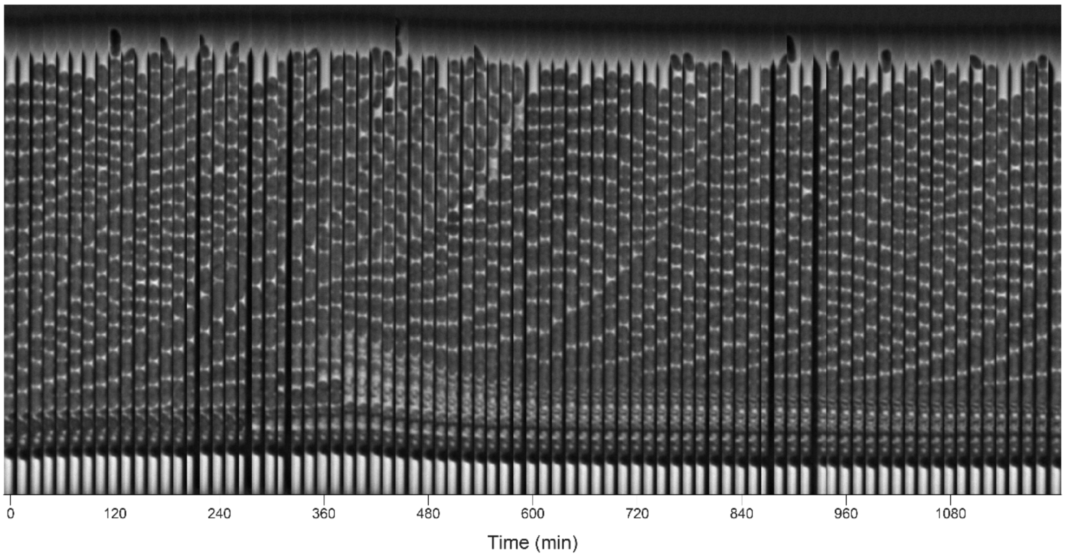


b

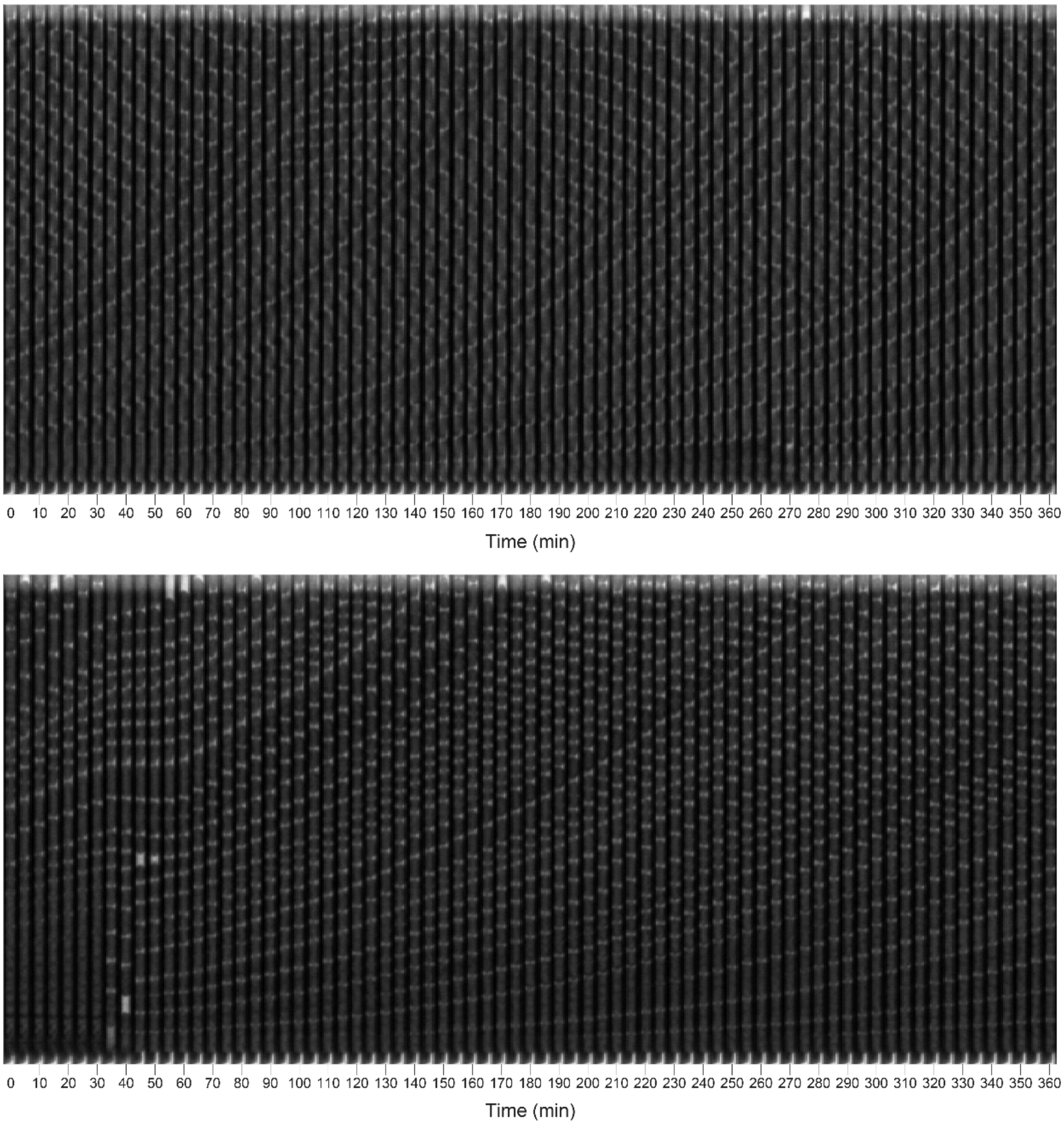


**Supplementary Fig. 7 | *galE* transposon mutant kymographs. a,** Trap from the pooled experiment.

The data is visualized in 15 min increments. **b,** Examples of two traps from the experiment with further investigation of the *galE* transposon mutants. The data is visualized in 5 min increments.

**Supplementary Video 1 |** Example video of a cell being picked from a trap and transferred to the flow line of the medium (tweezer) port. The video was recorded using Open Broadcast Software (OBS).

**Supplementary Table 1 |** List with the number of reads in each annotated gene from TIS on the whole Sucrose+ library (see separate file).

**Supplementary Table 2 |** List with the number of reads in each intergenic region from TIS on the whole Sucrose+ library (see separate file). The intergenic regions are named after their genomic position in the transposon library reference strain sequence (available on request).

**Supplementary Table 3 |** List containing the genes where transposons were mapped to across the pooled screens. Light blue: results were inconsistent between the pooled and arrayed screens; light yellow: results were consistent; white: not tested in an arrayed screen (see separate file).

**Supplementary Table 4 |** List of all strains (see separate file).

**Supplementary Table 5 |** List of all oligonucleotides (see separate file).

**Supplementary Table 6 |** Microscopy-related information for each panel. Includes the number of foci, number of traps, optical setup, microfluidic device type, trap size in said device, setting on the light source used for epi-fluorescence illumination, exposure time for said illumination, duration of the imaging experiment, concentration of F-108 Pluronic®, whether all colonies from a specific trap in the pooled transposon mutagenesis screens were PCR verified for their transposon location, and the number of replicates for each experiment (see separate file).
